# Multiple genetic variants at the *SLC30A8* locus affect local super-enhancer activity and influence pancreatic β-cell survival and function

**DOI:** 10.1101/2023.07.13.548906

**Authors:** Ming Hu, Innah Kim, Ignasi Morán, Weicong Peng, Orien Sun, Amélie Bonnefond, Amna Khamis, Silvia Bonas-Guarch, Philippe Froguel, Guy A. Rutter

## Abstract

Variants at the *SLC30A8* locus are associated with type 2 diabetes (T2D) risk. The lead variant, rs13266634, encodes an amino acid change, Arg325Trp (R325W), at the C-terminus of the secretory granule-enriched zinc transporter, ZnT8. Although this protein-coding variant was previously thought to be the sole driver of T2D risk at this locus, recent studies have provided evidence for lowered expression of *SLC30A8* mRNA in protective allele carriers. In the present study, combined allele-specific expression (cASE) analysis in human islets revealed multiple variants that influence *SLC30A8* expression. Epigenomic mapping identified an islet-selective enhancer cluster at the *SLC30A8* locus, hosting multiple T2D risk and cASE associations, which is spatially associated with the *SLC30A8* promoter and additional neighbouring genes. Deletions of variant-bearing enhancer regions using CRISPR-Cas9 in human-derived EndoC-βH3 cells lowered the expression of *SLC30A8* and several neighbouring genes, and improved insulin secretion. Whilst down-regulation of *SLC30A8* had no effect on beta cell survival, loss of *UTP23*, *RAD21 or MED30* markedly reduced cell viability. Although eQTL or cASE analyses in human islets did not support the association between these additional genes and diabetes risk, the transcriptional regulator JQ1 lowered the expression of multiple genes at the *SLC30A8* locus and enhanced stimulated insulin secretion.

## Introduction

Type 2 Diabetes (T2D) affects more than 400 million people worldwide and the total number of people affected by diabetes is expected to increase to >600 million by 2045 (https://www.diabetesatlas.org). While cultural and environmental factors are important elements in disease risk, genetics also plays a major role. Thus, more than 200 genetic loci carrying >400 independent signals associated with T2D have been discovered [1–4], with the majority of these influencing insulin secretion [5–10]. Analyses of chromatin accessibility and expression quantitative trait (eQTL) loci in human islets have further increased our understanding of how variants are likely to exert their effects [7, 11–13]. These, alongside functional studies to determine the role of implicated gene products in pancreatic β cell function, have allowed us [14–17] to determine the likely mode(s) of action of variants at several T2D-associated *loci*.

The *SLC30A8* locus was one of the first to be identified by genome-wide association studies (GWAS) as being associated with T2D [18] and has attracted considerable interest ever since [19, 20]. The lead single nucleotide polymorphism (SNP) rs13266634 is located in exon 8 of *SLC30A8*, a gene expressed almost exclusively in the endocrine pancreas. Rs13266634 is a non-synonymous variant that alters the primary sequence of the secretory granule zinc transporter ZnT8 (SLC30A8) at the C-terminus of the protein, likely affecting monomer dimerization [21–23], and thus intrinsic Zn^2+^ transporter activity. Nevertheless, the mechanisms through which this change may affect T2D risk remain unclear. Firstly, structural predictions and functional studies exploring the impact of the common R325W variant on ZnT8 have been inconclusive. Evidence for both diminished [21] and increased [23] intrinsic zinc transporter activity of the protein encoded by the risk (R325) variant has been obtained using different approaches.

Extensive studies examining the impact of deletion or inactivation of *SLC30A8* in humans, mice and in cell lines have yielded contradictory results. Thus, several [21, 24], though not all [19, 25], studies in mice have demonstrated a requirement for *Slc30a8* for normal insulin granule biogenesis and Zn^2+^ accumulation, and for normal insulin secretion *in vivo* and/or *in vitro*. In contrast, rare loss-of-function *SLC30A8* variants are beneficial for human β-cell function, lowering T2D risk [26, 27]. Correspondingly, the deletion of *SLC30A8* from human beta cells enhances insulin secretion [27], whilst the p.Arg138* loss of function (LoF) variant protected against apoptosis at low extracellular Zn^2+^ concentrations [28].

GWAS studies [22] have also identified a cluster of non-coding genetic variants at *SLC30A8* locus (https://t2d.hugeamp.org) [31], whose contribution to disease risk relative to rs13266634 is unclear. Recent findings [12] have suggested that carriers of protective alleles at this locus display lowered *SLC30A8* transcript levels. Whether these actions are due to altered chromatin accessibility at this locus, and hence altered transcription, or are mediated by post-transcriptional events such as altered mRNA stability, has not been assessed. Likewise, the possible contribution of genes which may be co-ordinately regulated with *SLC30A8* at this locus has not been explored.

Here, we sought to determine whether rs13266634 or other variants affect the transcription of *SLC30A8* or neighbouring genes, with consequences for β cell function or survival.

## Results

### Impact of T2D variants on transcript levels in human islets

GWAS-identified variants at the 3’ end of *SLC30A8* are shown in Figure S1 (www.t2d.hugeamp.org). To determine whether the variants were associated with the expression of the *SLC30A8* gene as well as nearby genes in human islets, we first examined expression quantitative trail *loci* (eQTL) analyses using data from multiple sources. As shown in Table 1A-1E, no statistically significant associations were identified.

**Table 1A.** eQTL analysis data set from laser capture cohort [57].

| rsID | REF | ALT | Distance (Kb) | Nominal P | FDR | beta | Gene Symbol |
| --- | --- | --- | --- | --- | --- | --- | --- |
| rs3802177 | G | A | 326852 | $6.37 \times 10^{-3}$ | 0.442 | 0.25 | RAD21 |
| rs13266634 | C | T | 298121 | $6.37 \times 10^{-3}$ | 0.442 | 0.18 | MIR3610 /<br>RAD21-AS1 |
| rs4300038 | G | A | 359742 | $6.66 \times 10^{-3}$ | 0.449 | 0.26 | RAD21 |
| rs13266634 | C | T | 37216 | $7.88 \times 10^{-3}$ | 0.476 | 0.36 | SLC30A8 |
| rs9650069 | C | T | 345847 | $8.36 \times 10^{-3}$ | 0.485 | 0.25 | RAD21 |
| rs11558471 | A | G | 299071 | $8.93 \times 10^{-3}$ | 0.495 | 0.16 | MIR3610 /<br>RAD21-AS1 |
| rs3802177 | G | A | 37458 | $1.31 \times 10^{-2}$ | 0.555 | 0.33 | SLC30A8 |
| rs13266634 | C | T | 326610 | $1.45 \times 10^{-2}$ | 0.570 | 0.22 | RAD21 |
| rs35859536 | C | T | 333302 | $1.55 \times 10^{-2}$<br>0.0154994 | 0.585 | 0.22 | RAD21 |
| rs11558471 | A | G | 327560 | $1.58 \times 10^{-2}$ | 0.583 | 0.21 | RAD21 |
| rs11558471 | A | G | 406989 | $1.66 \times 10^{-2}$ | 0.591 | 0.11 | UTP23 |
| rs3802177 | G | A | 298363 | $1.7 \times 10^{-2}$ | 0.602 | 0.16 | MIR3610 /<br>RAD21-AS1 |
| rs13266634 | C | T | 400600 | $1.84 \times 10^{-2}$ | 0.606 | 0.18 | UTP23 |
| rs35859536 | C | T | 43908 | $2.36 \times 10^{-2}$ | 0.641 | 0.31 | SLC30A8 |
| rs9650069 | C | T | 317358 | $2.47 \times 10^{-2}$ | 0.648 | 0.15 | MIR3610 /<br>RAD21-AS1 |
| rs35859536 | C | T | 304813 | $2.54 \times 10^{-2}$ | 0.652 | 0.15 | MIR3610 /<br>RAD21-AS1 |
| rs4300038 | G | A | 331253 | $3.68 \times 10^{-2}$ | 0.703 | 0.14 | MIR3610 /<br>RAD21-AS1 |
| rs13266634 | C | T | 326609 | $4.02 \times 10^{-2}$ | 0.715 | 0.077 | RAD21 |
| rs11558471 | A | G | 38166 | $4.13 \times 10^{-2}$ | 0.718 | 0.26 | SLC30A8 |
| rs3802177 | G | A | 400842 | $4.15 \times 10^{-2}$ | 0.719 | 0.16 | UTP23 |
| rs16889471 | G | A | 303958 | $4.53 \times 10^{-2}$ | 0.730 | -0.15 | MIR3610 /<br>RAD21-AS1 |
| rs13266634 | C | T | -3724 | $4.90 \times 10^{-2}$ | 0.734 | 0.15 | SLC30A8 |

**Table 1B.** eQTL analysis of organ donors (beta cells) [52].

| rsID | REF | ALT | Distance (Kb) | Nominal P value | FDR | beta | Gene Symbol |
| --- | --- | --- | --- | --- | --- | --- | --- |
| rs16889471 | G | A | -933472 | $5.49 \times 10^{-3}$ | 0.927 | -0.68 | EXT1 |
| rs11774700 | T | C | 441106 | $1.74 \times 10^{-2}$ | 0.942 | -0.57 | EIF3H |
| rs11558471 | A | G | 298694 | $2.06 \times 10^{-2}$ | 0.945 | -0.83 | MIR3610 |
| rs13266634 | C | T | -939309 | $2.52 \times 10^{-2}$ | 0.949 | 0.61 | EXT1 |
| rs3802177 | G | A | -939067 | $2.52 \times 10^{-2}$ | 0.949 | 0.61 | EXT1 |
| rs4300038 | G | A | -906177 | $2.52 \times 10^{-2}$ | 0.949 | 0.61 | EXT1 |
| rs9650069 | C | T | -920072 | $2.52 \times 10^{-2}$ | 0.949 | 0.61 | EXT1 |
| rs11558471 | A | G | 406569 | $2.59 \times 10^{-2}$ | 0.945 | -0.50 | EIF3H |
| rs13266634 | C | T | 405619 | $2.66 \times 10^{-2}$ | 0.950 | -0.50 | EIF3H |
| rs3802177 | G | A | 405861 | $2.66 \times 10^{-2}$ | 0.950 | -0.50 | EIF3H |
| rs4300038 | G | A | 438751 | $2.66 \times 10^{-2}$ | 0.950 | -0.50 | EIF3H |
| rs9650069 | C | T | 424856 | $2.66 \times 10^{-2}$ | 0.950 | -0.50 | EIF3H |
| rs35859536 | C | T | 15268 | $2.71 \times 10^{-2}$ | 0.951 | 0.29 | RN7SL826P |
| rs35859536 | C | T | -932617 | $2.90 \times 10^{-2}$ | 0.950 | 0.63 | EXT1 |
| rs13266634 | C | T | -348169 | $3.09 \times 10^{-2}$ | 0.952 | 0.67 | MED30 |
| rs3802177 | G | A | -347927 | $3.09 \times 10^{-2}$ | 0.953 | 0.67 | MED30 |
| rs4300038 | G | A | -315037 | $3.09 \times 10^{-2}$ | 0.953 | 0.67 | MED30 |
| rs9650069 | C | T | -328932 | $3.09 \times 10^{-2}$ | 0.953 | 0.67 | MED30 |
| rs11558471 | A | G | -347219 | $3.27 \times 10^{-2}$ | 0.954 | 0.67 | MED30 |
| rs11774700 | T | C | 441106 | $3.57 \times 10^{-2}$ | 0.955 | -0.39 | EIF3H |
| rs13266634 | C | T | 8576 | $4.88 \times 10^{-2}$ | 0.960 | 0.25 | RN7SL826P |
| rs3802177 | G | A | 8818 | $4.88 \times 10^{-2}$ | 0.960 | 0.25 | RN7SL826P |
| rs4300038 | G | A | 41708 | $4.88 \times 10^{-2}$ | 0.960 | 0.25 | RN7SL826P |
| rs9650069 | C | T | 27813 | $4.88 \times 10^{-2}$ | 0.960 | 0.25 | RN7SL826P |
| rs11774700 | T | C | -903822 | $4.98 \times 10^{-2}$ | 0.960 | 0.46 | EXT1 |
| rs11774700 | T | C | -903822 | $4.98 \times 10^{-2}$ | 0.960 | -0.51 | EXT1 |

**Table 1C.** eQTL analysis from TIGER cohort [12].

| rsID | EA | NEA | Gene | Nominal P | FDR | Effect (Z) |
| --- | --- | --- | --- | --- | --- | --- |
| rs13266634 | C | T | SLC30A8 | 0.367 | NS | 0.903 |
| rs13266634 | C | T | MED30 | 0.979 | NS | 0.026 |
| rs13266634 | C | T | EXT1 | 0.952 | NS | 0.06 |
| rs13266634 | C | T | RAD21-AS1 | 0.784 | NS | 0.274 |
| rs3802177 | G | A | EXT1 | 0.938 | NS | 0.078 |
| rs3802177 | G | A | RAD21-AS1 | 0.873 | NS | 0.159 |
| rs35859536 | C | T | EXT1 | 0.873 | NS | -0.055 |
| rs35859536 | C | T | RAD21-AS1 | 0.961 | NS | -0.05 |
| rs11558471 | G | A | SLC30A8 | 0.594 | NS | -0.534 |
| rs11558471 | G | A | MED30 | 0.905 | NS | 0.119 |
| rs11558471 | G | A | Mir3610 | NA | NA | NA |
| rs11558471 | G | A | RAD21-AS1 | 0.886 | NS | -0.144 |
| rs11558471 | G | A | RP11-654G14.1 | 0.648 | NS | 0.457 |
| rs4300038 | G | A | MED30 | 0.892 | NS | -0.136 |
| rs4300038 | G | A | EXT1 | 0.863 | NS | -0.173 |
| rs4300038 | G | A | RAD21-AS1 | 0.949 | NS | -0.065 |
| rs9650069 | C | T | MED30 | 0.817 | NS | -0.232 |
| rs9650069 | C | T | RAD21-AS1 | 0.770 | NS | -0.293 |
| rs9650069 | C | T | EXT1 | 0.973 | NS | -0.034 |
| rs11774700 | C | T | MED30 | 0.562 | NS | -0.58 |
| rs11774700 | C | T | RAD21-AS1 | 0.791 | NS | 0.265 |
| rs11774700 | C | T | EXT1 | 0.538 | NS | 0.616 |

**Table 1D.** eQTL analysis from InsPIRE cohort [64].

| SNP | EA | NEA | Gene | P value | FDR | Effect (Z) |
| --- | --- | --- | --- | --- | --- | --- |
| rs13266634 | T | C | SLC30A8 | 0.390 | NA | -0.016398 |
| rs13266634 | T | C | MED30 | 0.038 | NS | -0.058180 |
| rs13266634 | T | C | EXT1 | 0.603 | NA | 0.0124346 |
| rs13266634 | T | C | RAD21-AS1 | NA | NS | NA |
| rs3802177 | A | G | EXT1 | 0.625 | NA | 0.0116709 |
| rs3802177 | A | G | RAD21-AS1 | NA | NA | NA |
| rs35859536 | T | C | EXT1 | 0.567 | NA | 0.0136128 |
| rs35859536 | T | C | RAD21-AS1 | NA | NA | NA |
| rs11558471 | G | A | SLC30A8 | 0.361 | NA | -0.0172161 |
| rs11558471 | G | A | MED30 | 0.024 | NA | -0.0624528 |
| rs11558471 | G | A | MIR3610 | NA | NA | NA |
| rs11558471 | G | A | RAD21-AS1 | NA | NA | NA |
| rs11558471 | G | A | RP11-654G14.1 | 0.488 | NA | -0.025098 |
| rs4300038 | A | G | MED30 | 0.046 | NA | -0.055691 |
| rs4300038 | A | G | EXT1 | 0.592 | NA | 0.0127821 |
| rs4300038 | A | G | RAD21-AS1 | NA | NA | NA |
| rs9650069 | T | C | MED30 | 0.032 | NA | -0.059416 |
| rs9650069 | T | C | RAD21-AS1 | NA | NA | NA |
| rs9650069 | T | C | EXT1 | 0.695 | NA | 0.00933485 |
| rs11774700 | C | T | MED30 | 0.025 | NA | -0.0641155 |
| rs11774700 | C | T | RAD21-AS1 | NA | NA | NA |
| rs11774700 | C | T | EXT1 | 0.276 | NA | 0.0266843 |

EA = effect allele, NEA = non-effect allele

**Table 1E.** eQTL analysis from Atla et al [53].

| rsID | EA | NEA | Gene | Nominal P | FDR | Effect (Z) |
| --- | --- | --- | --- | --- | --- | --- |
| rs13266634 | T | C | SLC30A8 | 0.268 | NS | -0.0394087 |
| rs13266634 | T | C | MED30 | 0.539 | NS | -0.0226174 |
| rs13266634 | T | C | EXT1 | NA | NA | NA |
| rs13266634 | T | C | RAD21-AS1 | NA | NA | NA |
| rs3802177 | A | G | EXT1 | NA | NA | NA |
| rs3802177 | A | G | RAD21-AS1 | NA | NA | NA |
| rs35859536 | T | C | EXT1 | NA | NA | NA |
| rs35859536 | T | C | RAD21-AS1 | NA | NA | NA |
| rs11558471 | G | A | SLC30A8 | 0.211 | NS | -0.0439515 |
| rs11558471 | G | A | MED30 | 0.686 | NS | -0.0439515 |
| rs11558471 | G | A | MIR3610 | NA | NA | NA |
| rs11558471 | G | A | RAD21-AS1 | NA | NA | NA |
| rs11558471 | G | A | RP11-654G14.1 | 0.162 | NS | -0.0717920 |
| rs4300038 | G | A | MED30 | 0.562 | NS | -0.0717920 |
| rs4300038 | G | A | EXT1 | NA | NA | NA |
| rs4300038 | G | A | RAD21-AS1 | NA | NA | NA |
| rs9650069 | T | C | MED30 | 0.662 | NA | -0.0162419 |
| rs9650069 | T | C | RAD21-AS1 | NA | NA | NA |
| rs9650069 | T | C | EXT1 | NA | NA | NA |
| rs11774700 | C | T | MED30 | NA | NA | NA |
| rs11774700 | C | T | RAD21-AS1 | NA | NA | NA |
| rs11774700 | C | T | EXT1 | NA | NA | NA |

EA = effect allele, NEA = non-effect allele

Given the absence of eQTL associations we turned next to combined allele-specific expression (cASE) analyses, interrogating published RNA-seq data from heterozygous islet samples [12]. Because both alleles in heterozygotes are exposed to the same milieu, this approach circumvents confounding effects of non-genetic variables (cold ischemia time, donor health and demographic variables, pre-mortem therapies, etc). Figure 1A illustrates the location of the four significant cASE reporter variants in the *SLC30A8* gene.

**Figure 1.**
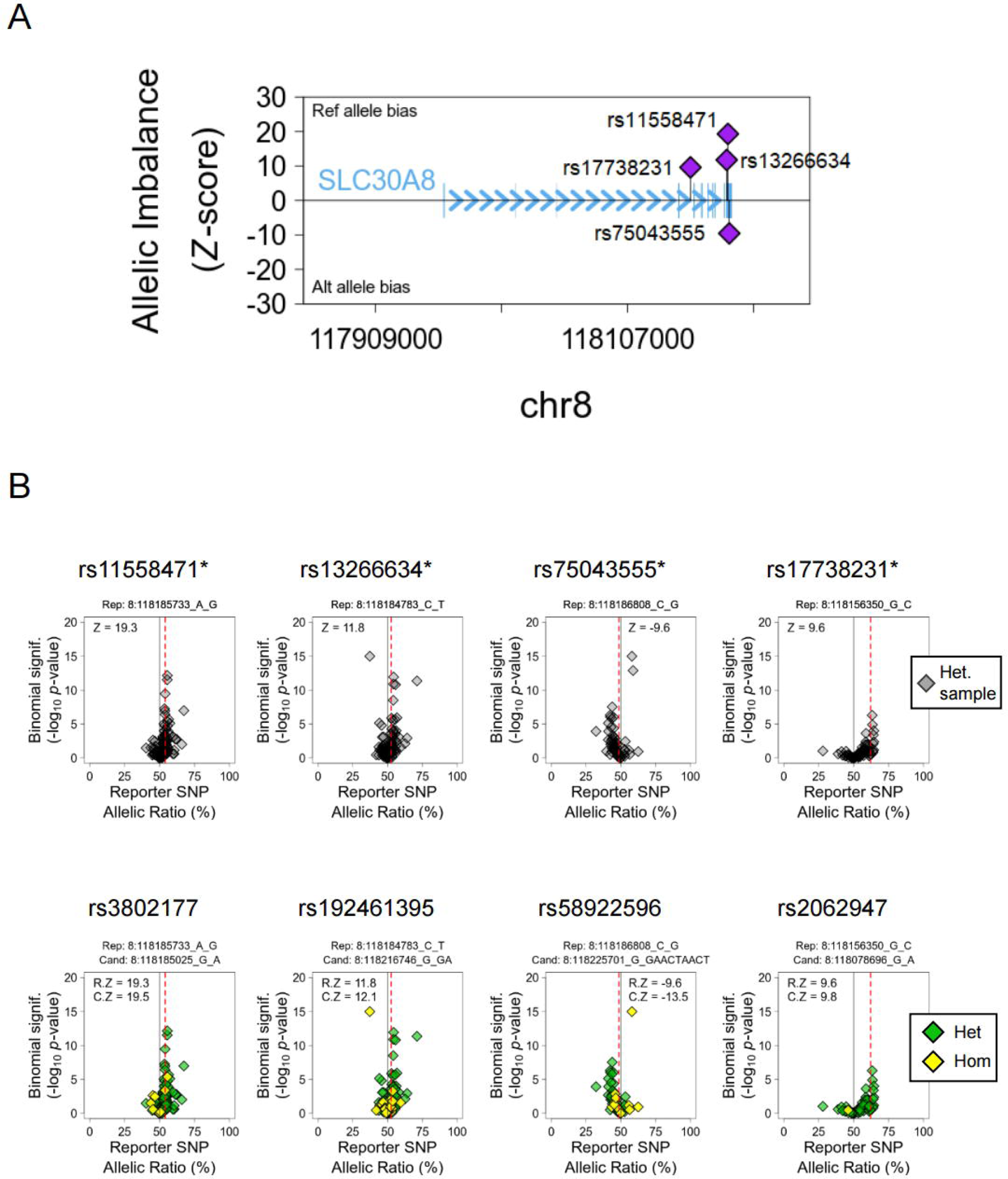
Combined Allele-Specific Expression (cASE) Analysis at the *SLC30A8* locus. A. Location of the reporter variants with significant cASE signal. The y-axis shows the strength and direction of their effect (Z-score). B. Detail of *SLC30A8* reporter variants (top panel) and their top candidate cis-regulatory variant to drive the cASE effect (bottom panel). The samples are grouped by the genotype of the candidate variant, heterozygous (Het., green), and homozygous (Hom., yellow). The first group needs to have a significant cASE for the gene (C.Z), while the second need to show no significant cASE. If C.Z is stronger than the reporter cASE calculated with all the samples (R.Z), the variant is considered a candidate.

We extended our earlier cASE analysis at this locus [12], demonstrating that highly correlated rs11558471 and rs13266634 variants (LD r^2^ = 0.96) exhibited a strong allelic bias of *SLC30A8* expression (*p*<4.64 x 10^-14^ and *p*<2.88 x 10^-6^ respectively) (Figure 1B, upper panel). The rs13266634 T2D risk allele (C) is the preferential allele for transcription. Additionally, we identified two independent cASE signals, rs17738231 (*p*<1.06 x 10^-4^) and rs75043555 (*p*<1.13 x 10^-4^), which have no known GWAS associations (Table 2). Candidate regulatory variants mediating the *SLC30A8* cASE signal are reported in Figure 1B (bottom panel). In sum, these data indicate that the allele-biased expression of *SLC30A8* overlapped with previous GWAS signals, and identified independent non-GWAS variants that may also play a significant role in the transcriptional regulation of the gene.

**Table 2.** cASE analysis of heterozygous human islet samples [12].

| rsID | EA | NEA | Gene | P value | Z-score | Tag SNP | LD r2 |
| --- | --- | --- | --- | --- | --- | --- | --- |
| rs11558471 | A | G | SLC30A8 | $4.64 \times 10^{-14}$ | 19.30 | NA | NA |
| rs13266634 | C | T | SLC30A8 | $2.88 \times 10^{-6}$ | 11.75 | rs11558471 | 0.96 |
| rs17738231 | G | C | SLC30A8 | $1.06 \times 10^{-4}$ | 9.60 | rs11558471 | 0.06 |
| rs75043555 | C | G | SLC30A8 | $1.13 \times 10^{-4}$ | -9.56 | rs11558471 | 0.05 |

EA = effect allele, NEA = non-effect allele

Using the same strategy, we performed cASE analysis to detect differential expression of nearby genes. No significant islet cASE associations were apparent for *RAD21, MED30*, EXT1, *UTP23* or *RP11-654G14.1. RAD21-AS1* could not be examined due to the absence of suitable variants in the transcript (data not shown). Overall, these data suggest that the variants located in the region are associated with allele specific expression of *SLC30A8* but not of nearby genes.

To explore the molecular mechanisms that elicit allele-specific expression of *SLC30A8,* and to determine whether altered transcription was likely to contribute to T2D risk, we turned to local epigenomic maps of human islets (http://epigenomegateway.wustl.edu/browser/). These maps identified a high occupancy cluster of active enhancers [32] (Figure 2A, S2 and S3A). This cluster, stretching about ∼293 kb, contains several super-enhancer elements and is characterized by high levels of histone-H3 lysine-27 acetylation (H3K27ac), binding of transcriptional regulator CCCTC binding factor CTCF, and cohesin complex (Figure 2A and S3A), as well as multiple islet specific-transcription factors (Figure 2A and S3B). Analyses of epigenomic maps in other human subjects (http://compbio2.mit.edu/epimap_vis/) [33] suggested that this enhancer-rich domain is chiefly active in pancreatic islets (Figure S2 and Supplementary Data 1). Accordingly, *SLC30A8* mRNA is almost exclusively expressed in islets in human (https://tiger.bsc.es) [12] and in other mammals [21, 24].

**Figure 2.**
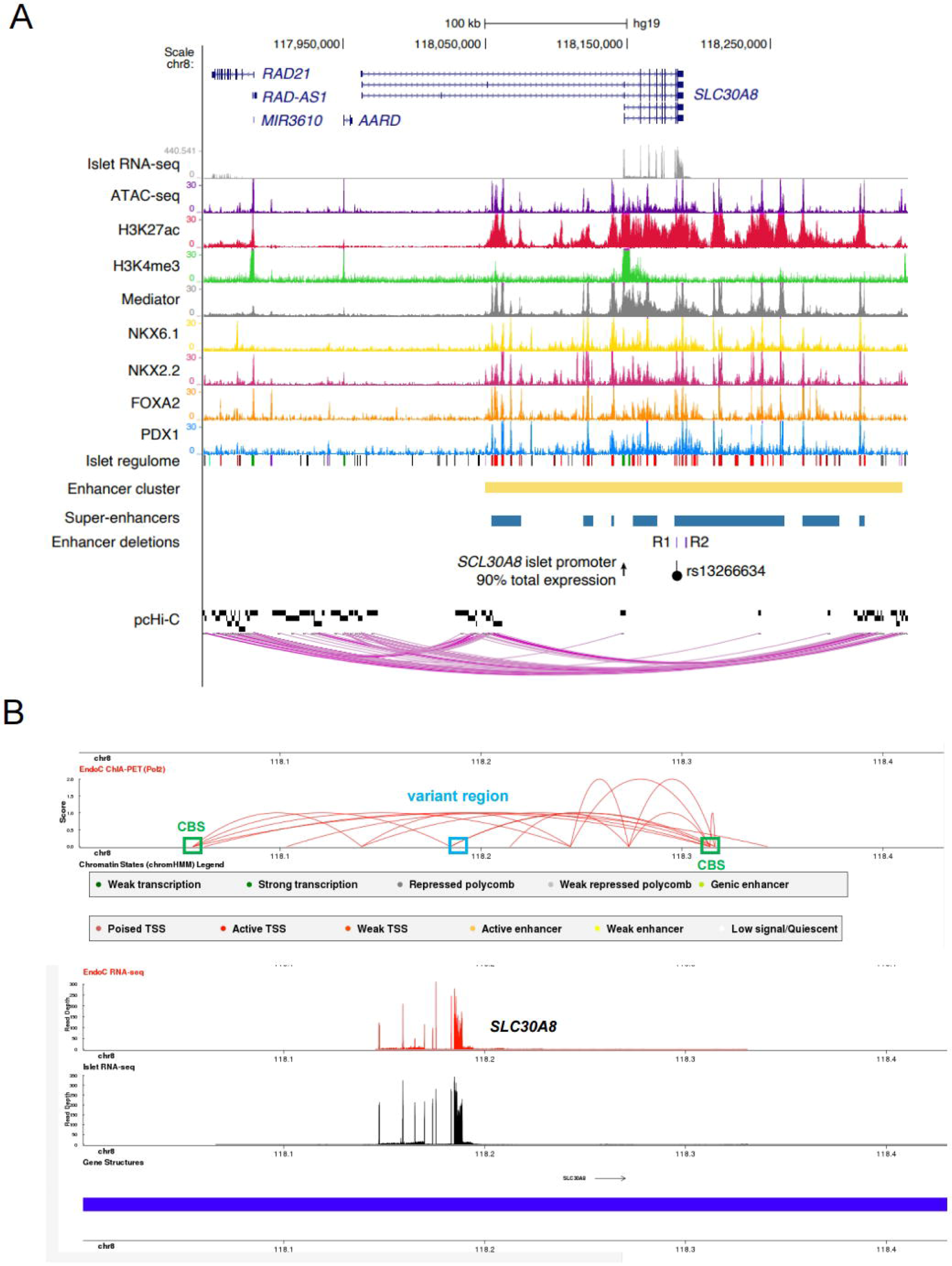
Combined epigenetic and chromatin maps at the *SLC30A8* locus. A. Integrative map of the *SLC30A8* locus in human islets showing chromatin state profiles and chromatin interactions between the super-enhancer (in blue) and *SLC30A8* and nearby genes, as determined by pcHi-C. R1 and R2 enhancer elements in the super-enhancer domain are coloured in purple. The lead T2D GWAS variant rs13266634 is represented as a black dot. B. CHIA-PET map [65] of the *SLC30A8* locus showing spatial contacts between the variant region and nearby regions. Polymerase II antibody was used to pull down protein/genomic DNA complex followed by whole genome sequencing [65]. Blue box: the variant region. Green box: CTCF binding sites (CBS) representing the 5’ and 3’ end of the super-enhancer cluster. Note that the variant region is spatially associated with the CTCF binding sites of the enhancer cluster.

Hi-C [34], Promoter-HiC [11] and CHIA-PET [35] analyses of human islet samples demonstrated that a group of genes at the 5’ end of the *SLC30A8* locus, including *RAD21* and *RAD21-AS1,* is spatially associated with the enhancer rich region through chromatin loop formation (Figure 2A and S3A). T2D risk and cASE associations are embedded within a single super-enhancer element nearby the *SLC30A8* promoter, suggesting that these transcripts may be co-regulated by the variant-bearing super-enhancer region (Figure 2A and S3A-B). These maps indicate that within the same topologically associating domain (TAD), the islet super-enhancer and embedded GWAS variants might influence the expression levels of not only *SLC30A8* but also *UTP23*, *RAD21*, *MED30* and *EXT1* genes and two long noncoding RNAs *RAD21-AS1* and *RP11-654G14.1*.

Several of the above genes encode components of complexes involved in fundamental cellular processes. *RAD21* is a cleavable component of the cohesin complex, involved in chromosome segregation, DNA replication, and other events [36]. The cohesin complex forms a ring structure that allows DNA to be extruded, controlling DNA looping. *UTP23* is a component of the small subunit processome (SSU) involved in rRNA-processing, ribosome biogenesis and 18S rRNA maturation [37]. *MED30* (mediator of RNA polymerase II transcriptional subunit 30) is part of the mediator complex bridging the enhancers and promoters [38]. *EXT1*, which encodes Exostosin-1 protein, is one of the two endoplasmic reticulum-resident type II transmembrane glycosyltransferase. Loss of function mutations in *EXT1* affect L-cell mass and insulin secretion in humans [39]. *RAD21-AS1* is expressed in the opposite direction to *RAD21*, while *RP11-654G14.1* is located in an intron within *SLC30A8*. Their roles in β-cells are unknown, and may include actions as enhancer RNAs (eRNA) [40].

### The super-enhancer regulates the expression of multiple genes within the same TAD domain

CTCF plays an important role in the formation of higher-order chromatin structures and may act as an insulator or boundary between *cis*-regulatory elements and their target genes [41, 42]. As shown in promoter-HiC map (Figure S3A), this enhancer-rich domain loops through CTCF binding sites (CBSs) at both 5’ and 3’ ends. With the guidance of epigenomic maps as above, we next attempted to explore the transcriptional role of the T2D and cASE variant-bearing super-enhancer region by determining whether multiple gene(s) were under its control.

We mutated the adjacent CBS sites individually or in combination. First, we mutated five CBS sites individually through CRISPR-Cas9 genome editing (Figure 3A and 3B, Figure S4A). The impact of these changes was assessed by two methods: 1. CTCF immunoprecipitation using CTCF antibody followed by qPCR. As shown in Figure 3C and S4B, CTCF binding was dramatically reduced after genome editing in all five CBS-mut cells. 2. We examined the CBS mutation efficiencies using SYBR green qPCR analysis. The mutation efficiencies were high (60-85%, Figure S4C), corroborating the results from the CTCF-pull-down assay.

**Figure 3:**
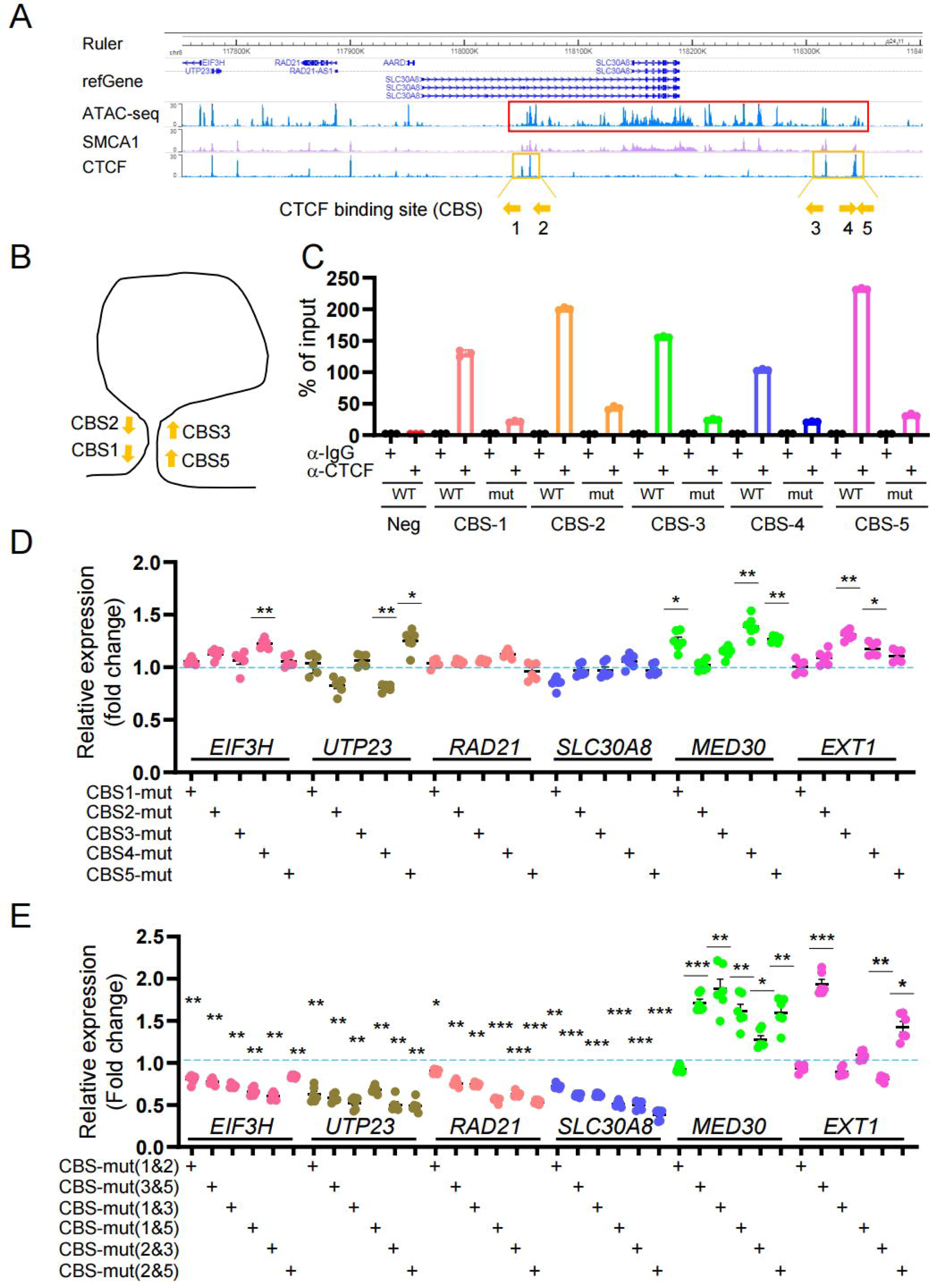
Disruption of the super-enhancer region by loss of CTCF binding impacts multiple genes. A. Location of CTCF binding site (CBS) at the 5’ and 3’ end of the super enhancer. Red box: Super-enhancer; Orange box and line: CTCF binding area. Orange arrow: Orientation of CFCT-binding sites. B. Diagram of convergent CTCF-binding at the base of loops. CTCF-binding sites 1 or 2 are potentially bind to 3 or 5 with high efficiency due to convergent orientation. The divergent orientation potentially formed between 1 or 2 and 4 is dis-favoured. Hence, 4 was excluded in the carton and not studied here for double mutation of CTCF binding site. C. CHIP-qPCR analysis for CTCF binding in wild type and CBS-mut cells. DD. qRT-PCR analysis of gene expression in single CBS-mut cells. Scramble gRNA infected cells were used as a control. Data are mean ± SEM. *, *P* < 0.05; **, *P* < 0.01; ***, *P* < 0.005. *n* = 3. E. qRT-PCR analysis of gene expression in double CBS-mut cells. Data are mean ± SEM. *, *P* < 0.05; **, *P* < 0.01; ***, *P* < 0.005. *n* = 3.

We next performed Taqman qRT-PCR analysis to examine the expression of genes in the TAD (Figure 3D). Whilst expression of *SLC30A8* and *RAD21* were unaltered, *EIF3H* and *UTP23*, which are located at the 5’ end of the super-enhancer domain, displayed minor changes compared with Scrambled control (set at 1.0) and we observed a trend of increased gene expression of *MED30* and *EXT1* (3’ of the super-enhancer). Since there are two to three CTCF binding sites on both 5’ and 3’ ends of this region, mutation of one of the CBS sites might not be sufficient to alter local chromatin organization, thus transcription. Hence, we next attempted to simultaneously mutate two CBS sites in order to alter the entire chromatin structure (Figure 3B). Mutation efficiencies achieved similar rates to single ones (Figure S4D). Interestingly, double mutations led to significant changes at the expression levels of all genes examined (Figure 3E): the genes at the 5’ end of the super-enhancer had significant low expression along with *SLC30A8*. *MED30* at the 3’ end showed significant transcriptional activity in 4 out of 6 combinations. The expression pattern of *EXT1* gene was also altered. Notably, mutations at CBS3&5 had significant changes on all genes, lowering the expression of genes at the 5’ end and increasing the expression at the 3’ end. Similar expression pattern was also observed in CBS-mut(2&5) cells. Likewise, the increased expression of *MED30* was associated with mutations of either CBS3 or CBS5 suggesting that these two CBSs are associated with enhancer elements regulating *MED30* expression. Taken together, these data suggest that the super-enhancer domain regulates the expression of not only *SLC30A8* but also nearby genes.

### T2D GWAS variants are located in the SLC30A8 super enhancer

Exploration of the credible set at this locus indicated that rs13266634 carries the highest posterior probability for T2D risk (Table 3). To determine whether any of the other variants may drive altered gene expression we performed a functional screen to assess their activity in human EndoC-βH3 cells. Five out of seven GWAS-identified variants were selected for these studies (Figure S1). Three variants (rs13266634, rs3802177, rs35859536) lie within two enhancer regions (termed R1, R2) located in two islet enhancer elements. Both R1 and R2 regions are located at the 3’ end of the *SLC30A8* locus and are physically close to the CBS sites as assessed by CHIA-PET mapping (Figure 2B) [34]. R1 overlaps with exon 8 of *SLC30A8* and hosts two variants, rs13266634, in the protein coding region, and rs3802177 in the 3’ UTR. *In silico* analysis of transcription factor binding using JASPAR [43] suggested differential transcription factor binding (Table 4) between T2D risk and protective variants. Two variants (rs3802177 and rs11558471) are located in the 3’ UTR of *SLC30A8* and may thus affect *SLC30A8* mRNA stability. *In silico* analysis suggests altered miRNA binding between the risk and protective variants (Table 4).

**Table 3.** Variant credible set at the *SLC30A8* locus.

| rsID | REF | ALT | Lead | MEGA_P | Posterior Probability | In credset? |
| --- | --- | --- | --- | --- | --- | --- |
| rs13266634 | C | T | rs13266634 | $3.22 \times 10^{-115}$ | 0.99830716 | YES |
| rs3802177 | G | A | rs13266634 | $7.67 \times 10^{-114}$ | 0.001647658 | NO |
| rs11558471 | A | G | rs13266634 | $4.38 \times 10^{-111}$ | $2.97 \times 10^{-6}$ | NO |
| rs35859536 | C | T | rs13266634 | $5.07 \times 10^{-113}$ | $4.22 \times 10^{-5}$ | NO |

**Table 4.** Candidate transcription factor and miRNA binding at variant sites.

| rsID | Risk variant | Protective variant |
| --- | --- | --- |
| rs13266634(C/T) | PAX9 | HIC2, ASCL1, NEUROD1 |
| rs3802177(G/A) | CREB1 | GATA6, LIN54, POU5F1 |
| rs35859536(C/T) | E2F6, FOXC1, HLTF, HMBOX1 | No |
| rs3802177(G/A) | No MiRNA binding | miR-190-5p or 190b-5p |
| rs11558471(A/G) | miR-1248 | miR-3074-5P |

R1 and R2 are deeply embedded in a single super-enhancer element located in the large enhancer cluster domain (Figure 2A and S3A). To assess whether R1 exhibits transcriptional activity, we cloned different lengths of the genomic DNA of the R1 region into a pGL4.32 reporter vector that carries a mini cytomegalovirus (CMV) promoter, and performed promoter-luciferase assays in EndoC-βH3 cells (Figure S5A-S5C). Luciferase activities of all DNA fragments were lower than empty vector indicating that this region is unlikely to have intrinsic enhancer activity, but may influence the activity of neighbouring enhancers.

To test this hypothesis in the context of genomic DNA, we deleted R1 from EndoC-H3 cells through CRISPR-Cas9-mediated genome editing [17]. Two gRNAs were designed to delete: (1) the entire R1 region (R1-del1) or (2), a small R1 region bearing two GWAS variants (R1-del2) (Figure 4A). Genomic DNA deletions were confirmed by Sanger sequencing (Figure S5D and S5E). The deletion efficiency was around 50-60% based on the protocol developed in our previous report (Figure S5F and S5G) [17]. Compared to control cells infected with scrambled gRNA sequences both R1-del1 and R1-del2 cells displayed lower expression of *SLC30A8* as well as several nearby genes (Figure 4B). Both R1-del1 and R1-del2 cells displayed increased insulin secretion stimulated by high glucose (15 vs 0.5 mM) and the phosphodiesterase inhibitor isobutylmethyl xanthine (IBMX; 0.5mM) (Figure 4C and 4D). Thus, lowered transcriptional activity brought about by R1-deletion improves β cell function.

**Figure 4.**
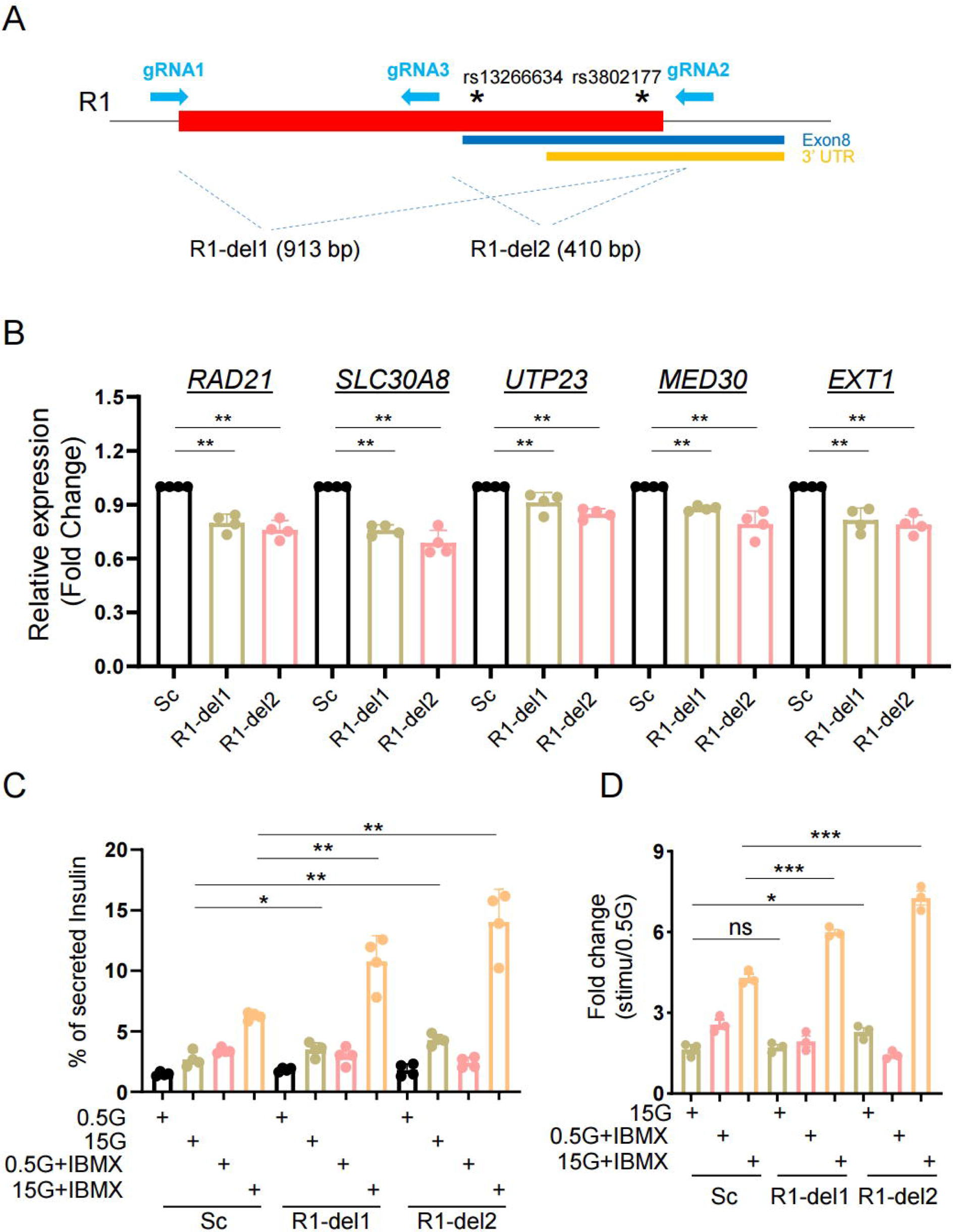
Role of R1 enhancer region in β-cell function. A. Diagram of R1 enhancer deletion in EndoC-βH3 by CRISPR-Cas9 mediated genome editing. Two gRNAs were designed to delete either entire R1 region (R1-del1 by gRNA1 and gRNA2) or a smaller region containing two variants (R1-del2 by gRNA3 and gRNA2). In both cases, deletions resulted into the loss of exon 8 of the *SLC30A8* gene, thus may affect ZnT8 translation. B. Taqman^TM^ qRT-PCR analysis of gene expression in R1-del cells. C. Glucose stimulated insulin secretion (GSIS) assay. Data are mean ± SEM. *, *P* < 0.05; **, *P* < 0.01; ***, *P* < 0.005. D. Fold change of secreted insulin. Data were normalized to insulin secretion at basal level (0.5 mM glucose). *n* = 3.

Whilst the above results may reflect an action of the genomic deletions on chromatin accessibility and transcription, it is also conceivable that the attendant change in the structure of the ZnT8 protein, notably through the deletion of part of the C-terminus (encoded by exon 8), may affect protein dimerization and hence secretory granule zinc uptake. This, in turn, may have consequences for regulated insulin secretion [19, 20]. We therefore examined the impact of manipulating the neighbouring R2 region, which bears rs35859536 (Figure 5A). This region lies downstream of the *SLC30A8* gene and, as such, its deletion does not affect the primary sequence of ZnT8 (Figure S3A). R2 showed lower transcriptional activity when compared with empty vector in promoter-luciferase assays (Figure S6A-S6C). However, the DNA fragment (F4) bearing rs35859536 had higher transcriptional activity than other regions within the R2 region (Figure S6B and S6C). Deletion of R2 (Figure 5A and Figure S6D-S6G) from EndoC-βH3 cells also lowered *SLC30A8* and nearby gene transcript levels, as observed after R1-deletion (Figure 5B). R2 deletion also enhanced insulin secretion stimulated by glucose and IBMX (Figure 5C and 5D). Taken together, these data indicate that both R1 and R2 participate in the transcriptional regulatory potential of the super-enhancer and impact the expression levels of *SLC30A8* and nearby genes.

**Figure 5.**
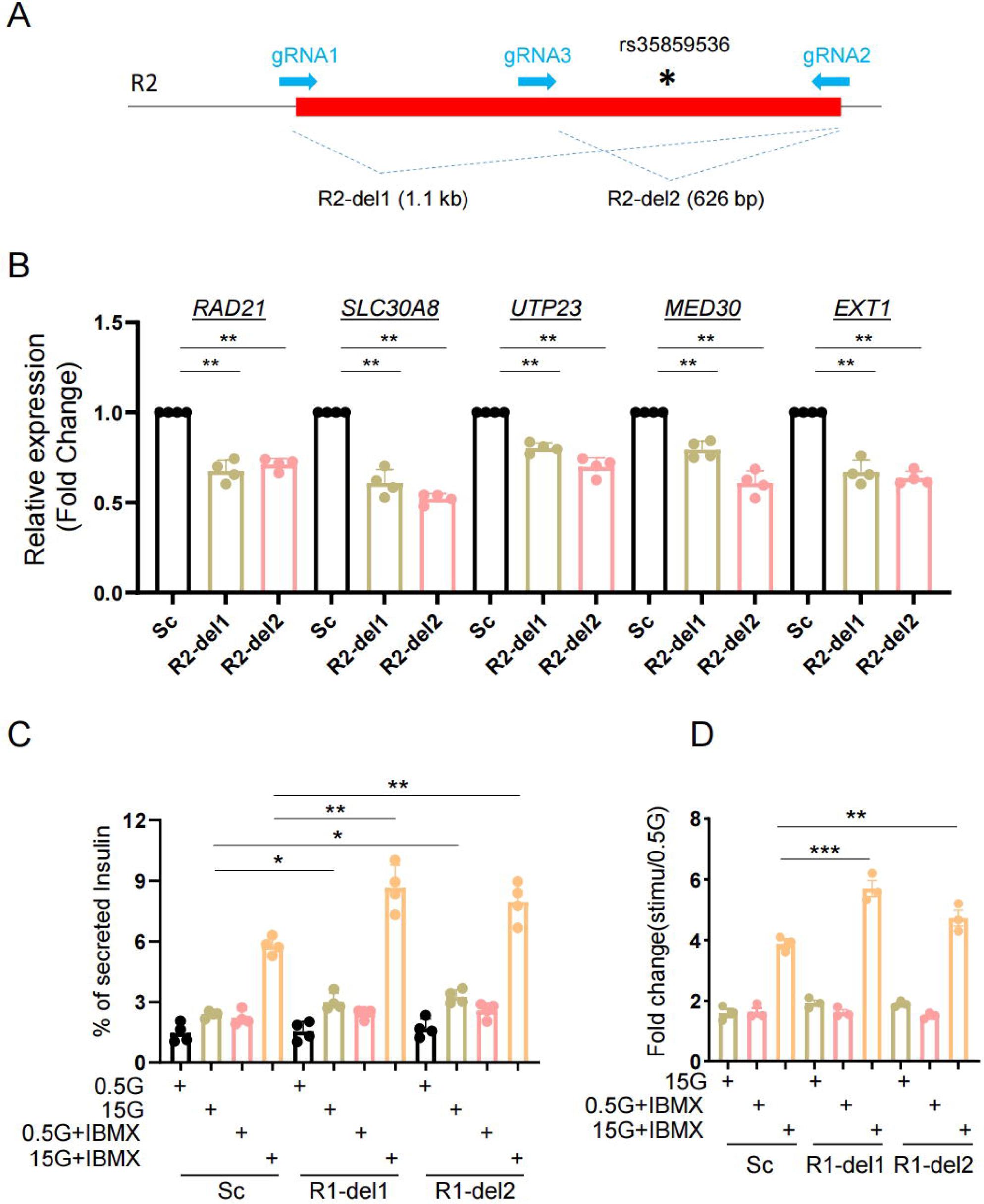
Role of R2 region in β-cell function. A. Diagram of R2 deletion in EndoC-βH3 by CRISPR-Cas9 mediated genome editing. Two gRNAs were designed to delete entire R2 region (R2-del1 by gRNA1 and gRNA2) or a region containing the variants (R2-del2 by gRNA3 and gRNA2). B. Taqman^TM^ qRT-PCR analysis of gene expression in R2-del cells. C. Glucose stimulated insulin secretion (GSIS) assay. Data are mean ± SEM. *, *P* < 0.05; **, *P* < 0.01; ***, *P* < 0.005. Fold change of secreted insulin. Data were normalized to insulin secretion at basal level (0.5 mM glucose). *n* = 3.

### T2D-associated variants affect transcriptional activity

We next investigated the roles of diabetes-associated variants in L cells. *In silico* analyses of transcription factor (TF) binding at the variant sites indicated that there may be differential TF binding (Table 4) which might affect the transcriptional activity of the active enhancers in which they reside. We first analysed the transcriptional activity of three variants in EndoC-LH3 cells (Figure 6A and 6B). The 200-300 bp DNA fragments surrounding either risk or protective alleles were cloned into the pGL4.32 vector (Figure 6A), which were confirmed by Sanger sequencing (Figure S7A). Promoter-luciferase assays in EndoC-βH3 cells (Figure 6B) revealed that the protective alleles for rs3802177 and rs35859536 displayed lower luciferase activity compared with their risk variants. No change (risk vs protective variant) was detected for rs13266634.

**Figure 6.**
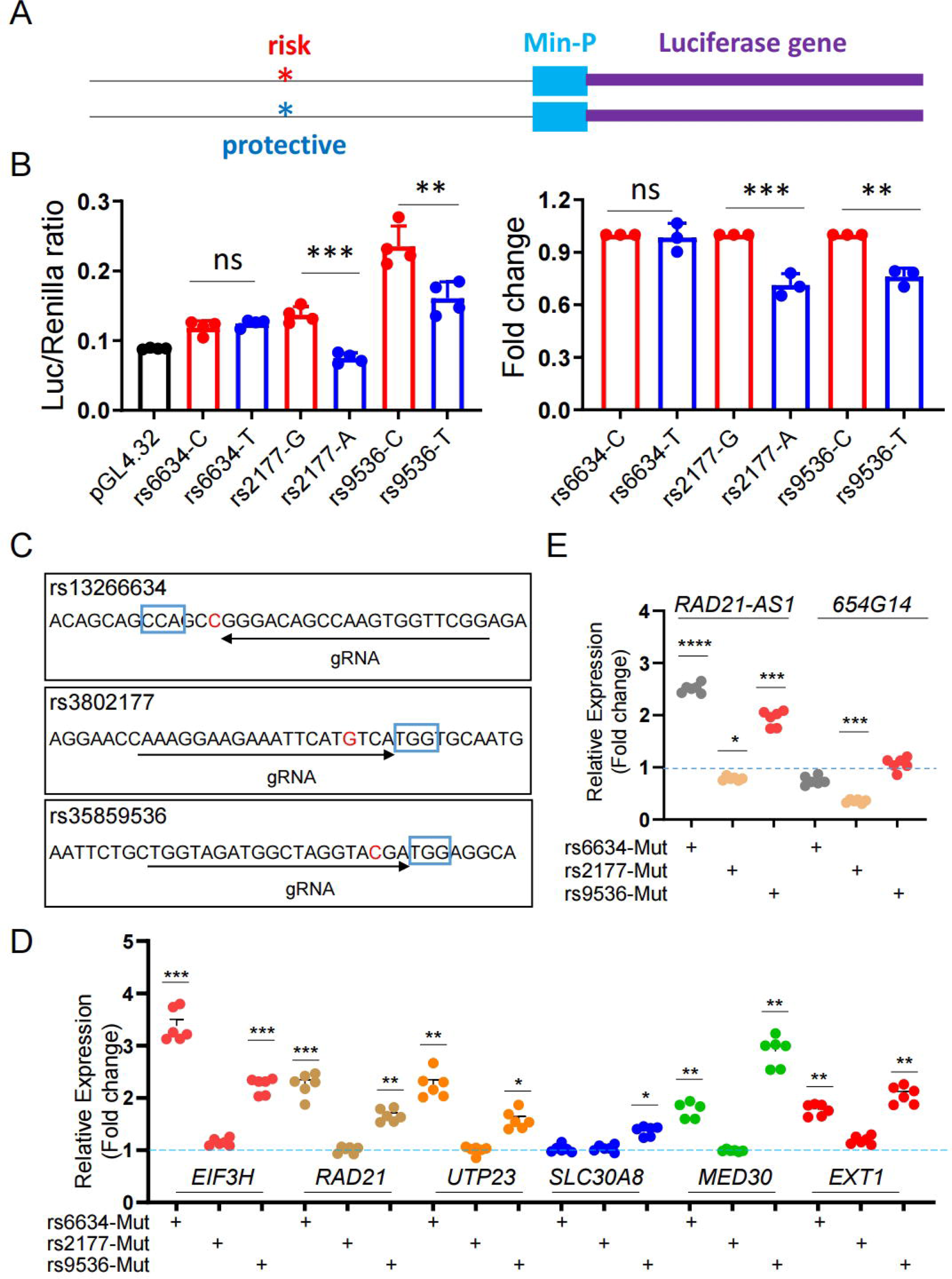
Function of individual variants in β-cells. A and B. Promoter-luciferase assay of individual variants in EndoC-βH3 cells. A. Diagram of variant bearing genomic DNA fragment in pGL4.23 vector. B. promoter-luciferase assay of the risk or protective variant in EndoC-βH3 cells. C. Diagram of blunt mutation at the variants by CRISPR-Cas9 mediated genome editing in EndoC-βH3 cells. D. Taqman^TM^ RT-qPCR analysis of *SLC30A8* and nearby genes. E. SYBR green RT-qPCR analysis of lncRNAs. Data are mean ± SEM. *, *P* < 0.05; **, *P* < 0.01; ***, *P* < 0.005. *n* = 3.

To determine whether these variants influence the expression of *SLC30A8* and / or nearby genes, we sought to introduce these variants into a L-cell line. However, both EndoC-βH1 and EndoC-βH3 cells are homozygous for the risk forms of the three variants and the introduction of the protective variants as “point mutations” by CRISPR-Cas9 engineering was not feasible in EndoC cell culture medium (Materials and Methods). To assess whether these risk variants may nevertheless have the potential to influence transcriptional activity, we created disruptive mutations at each variant site, with high efficiency, through conventional CRISPR-Cas9 genome editing (Figure 6C, S7B and S7C). Introduction of the mutations altered transcript levels at the *SLC30A8* locus as examined by TaqMan RT-qPCR for genes or SYBR-green RT-qPCR for IncRNAs (Figure 6D and 6E). Notably, disruption of rs13266634 led to higher expression not only of *SLC30A8* but also of other transcripts. Disruption of rs35839536 also caused significant changes at several genes, whereas rs3802177 disruption was without effect, though this may possibly be due to the lower efficiency of CRISPR-Cas9 genome editing at this site (Figure S7C).

### Genes within the TAD are required for cell survival

Based on chromatin accessibility and the presence of regulatory marks in human islets (Figure 2A and S3A), the *SLC30A8* super-enhancer is likely to co-regulate the *SLC30A8* gene as well as several nearby genes. Of these, the role of only *RAD21* [36] has previously been explored in β cells. We therefore inactivated each gene individually in EndoC-βH3 cells using CRISPR-Cas9 (Figure 7A and 7B, Figure S8A-S8C and Figure S9A-S9C). Inactivation of *RAD21* and *UTP23* resulted in a significant reduction of cell viability when compared with scrambled gRNA-treated control cells (Figure 7A and 7B; Figure S9C). *MED30* inactivation had a similar impact, although the effects on cell viability observed using gRNA2 were less marked than with the other guide RNAs deployed (Figure 7A and 7B, Figure S9C). In comparison, cells null for *SLC30A8* or *EXT1* showed no apparent defects in survival or growth (Figure 7A and 7B, Figure S9A-S9C).

**Figure 7.**
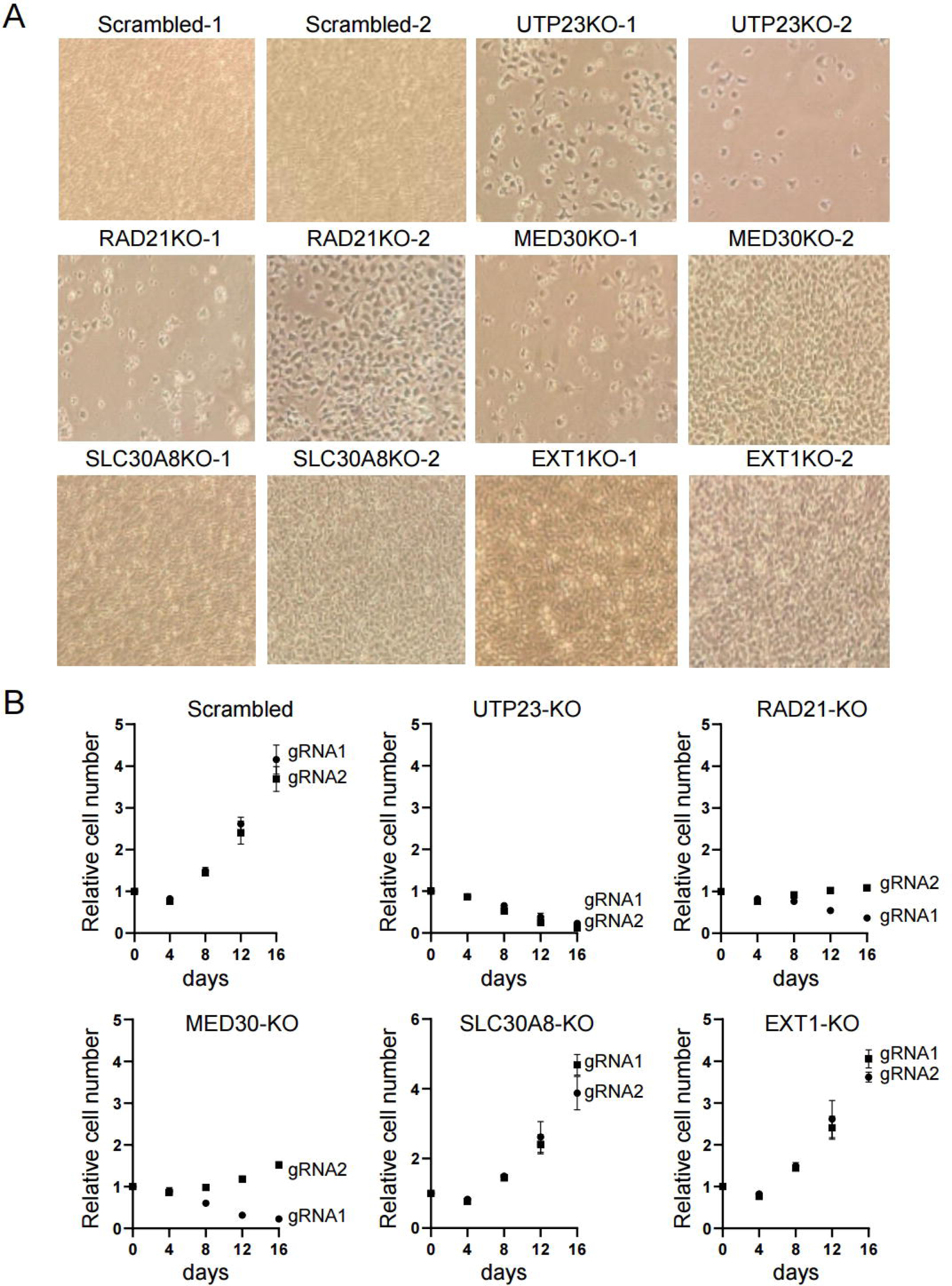
Impact of gene inactivation on cell survival. Two gRNAs were designed for each gene, cloned individually into pLenti-RIP-Cas9-BSD vector and delivered into 1 million EndoC-βH3 cells via lentiviral approach. Cells were selected with Blasticidin (25 μg/ml final concentration) for a week and cultured continuously for another 3 weeks. A. Morphologies of gene knockout cells. Photos were taken at the end of 4-week culture. Note that cells with scrambled gRNAs were confluent, along with *SLC30A8-KO* and *EXT1-KO* cells while *RAD21*, *MED30* and *UTP23* had significantly fewer surviving cells. B. Growth curves of control (scrambled) and gene-inactivated cells. At each time points, cells were fixed with formaldehyde and stained with crystal violet. Cell-associated dye was extracted with acetic acid and measured at 590 nm. The values were then normalized to the density at day 0.

### Inhibition of transcriptional activity by JQ1 improves β cell function

Our results above suggest that T2D risk variants at the *SLC30A8* locus increase the transcriptional activity of the enhancer-rich domain. Similar observations were also made in several GWAS-identified genetic loci where risk variants increase gene expression [7, 44, 45]. JQ1 is a potent inhibitor of Bromodomain and Extra-Terminal (BET) proteins. The BET protein family members such as BRD2, BRD3, BRD4, and BRDT are important regulators of epigenetic modifications and gene transcription [46]. Among these, BRD4 occupies super-enhancer regions and regulates gene transcription by binding to H3K27ac [47]. BRD4 inhibition disrupts the communication between SEs and their target promoters, and significantly decreases gene expression [47, 48]. JQ1 has been shown to increase insulin content and secretion in Rat INS1 (832/13) cells [49]. Based on these observations, we therefore sought to investigate whether JQ1 could reduce the transcriptional activity of the *SLC30A8* super-enhancer and in turn improve β cell function.

Examined in EndoC-βH3 and INS1 (832/13) cells, JQ1 increased insulin content in a dose dependent manner (Figure 8A and 8B; Figure S10A and S10B). we found that gene expression was altered mainly at higher doses of JQ1 (200 nM and 400 nM) with reduced levels for *EIF3H*, *SLC30A8* and *MED30* but higher level for *RAD21* in EndoC-βH3 cells (Figure 8C). Alterations of gene expression were more prominent in INS1 (832/13) cells in which *Slc30a8* expression was downregulated following the increase of JQ1 concentrations, along with *Eif3h*, *Utp23* at 50 nM and 100nM. *Med30* and *Ext1* were significantly increased but no change for *Rad1* (Figure S10C) was observed. To determine whether JQ1 influences β cell function we performed GSIS assays in the presence of high glucose (15 mM) or high glucose plus IBMX. As shown in Figure 8D and 8E, insulin secretion was slightly increased by JQ1 in the presence of high glucose and IBMX. These data are consistent with previous findings in INS1 (832/13) cells [49].

**Figure 8.**
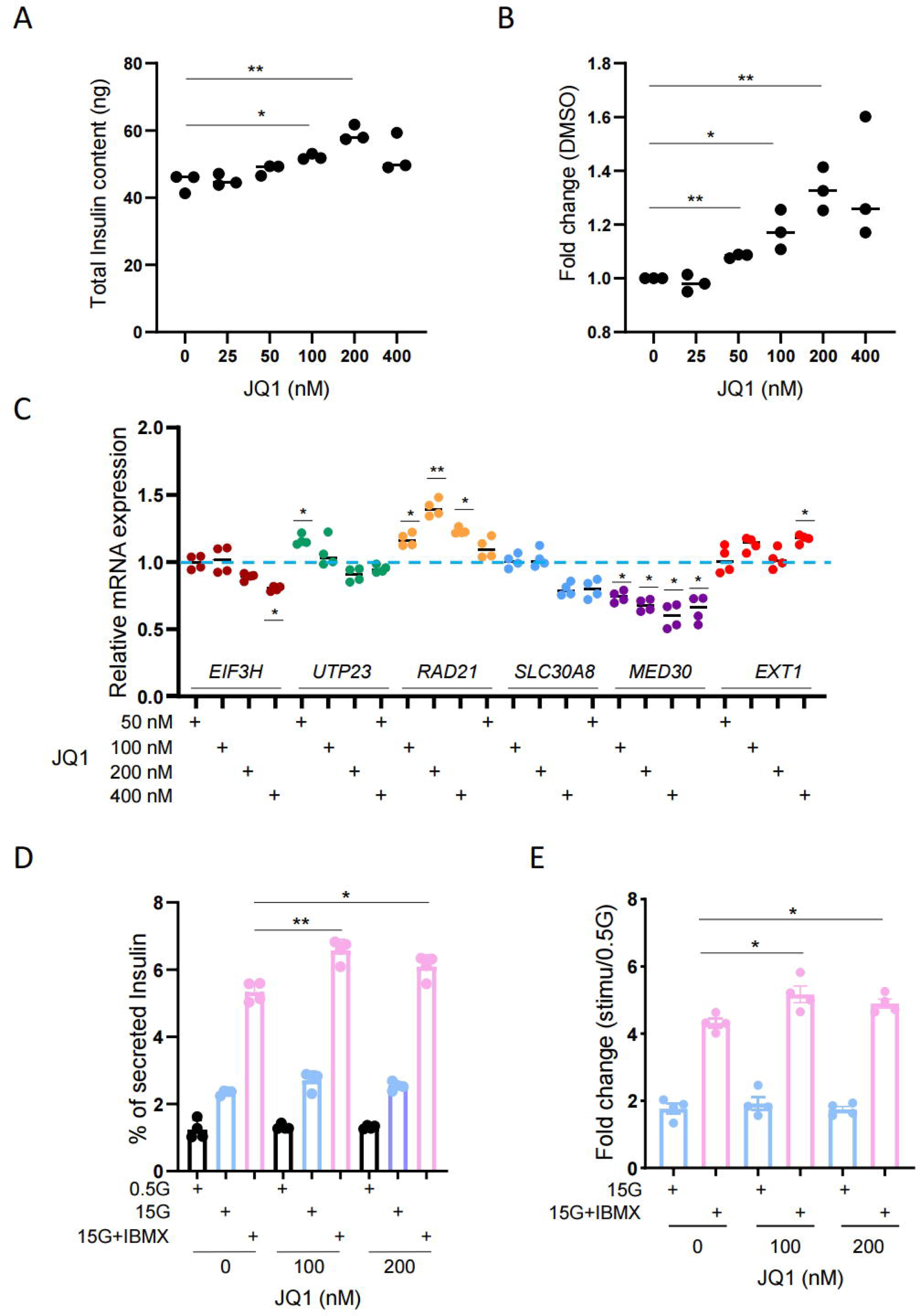
JQ1 increases insulin production and secretion in insulin producing cells. A. Total INSULIN content in JQ1 treated EndoC-βH3 cells. 7 x 10^4^ EndoC-βH3 cells were seeded in 96-well plate and incubated with different concentrations of JQ1 for 4 days. B. Fold change. DMSO treated sample was used as a control. C. Taqman^TM^ RT-qPCR analysis of *SLC30A8* and nearby genes in JQ1-treated EndoC-βH3 cells. EndoC-βH3 cells were treated with Tamoxifen (5 μm) for 3 weeks to remove gene cassette and increase insulin production [59]. Then, 1 x 10^6^ cells were seeded in 6-well plate and incubated with JQ1 for 4 days. D and E. Glucose induced insulin secretion (GSIS) assay in EndoC-βH3 cells. 7 x 10^4^ Tamoxifen treated cells were seeded in 96-well plate and incubated with JQ1 for 4 days. Cells were than preincubated with 2.8 mM glucose culture medium overnight before GSIS assay was performed. All Data above are mean ± SEM. *, *P* < 0.05; **, *P* < 0.01; ***, *P* < 0.005. *n* = 3.

## Discussion

### GWAS-identified genetic variants affect the expression of the SLC30A8 gene

In this study, we first assessed the effects of T2D GWAS variants on *SLC30A8* gene expression in human islet samples. Previous reports [20, 23, 24, 50] have focused almost exclusively on the lead variant rs13266634 which alters the amino acid sequence of ZnT8 (R325W), or on rare loss-of function *SLC30A8* variants [27, 51]. Whilst previous eQTL analyses [12, 52, 53] have failed to identify eQTLs for *SLC30A8*, analyses based on cASE identified significant allelic expression imbalance in *SLC30A8* [12]. Building on these earlier analyses [12], we confirm here that *SLC30A8* expression in human islets exhibits imbalanced allelic expression, where the T2D risk allele is more strongly expressed than the protective allele (Figure 1B). Among four reporter variants found in the *SLC30A8* transcript, rs11558471 displayed the most significantly imbalanced expression (*p*=4.64 x 10^-14^).

In contrast, we were unable to identify cASE of nearby genes when explored in the same collection of human islet samples [12]. Unlike *SLC30A8,* whose expression is restricted to the islet endocrine compartment [20], all neighbouring genes are expressed ubiquitously across tissues, including those co-isolated with human islets (ductal, acinar, mesenchymal etc. cells). Since the latter cell types comprise as much as 65 % of typical human islet preparations [53] our current study may be underpowered to detect cASE in the genes neighbouring *SLC30A8*. Future collective efforts to increase the scale of single-cell RNA-seq datasets facilitate cASE analysis at single β-cell resolution and they might expand the number of GWAS loci with allelic imbalanced gene expression.

### The SLC30A8 super-enhancer may regulate multiple genes

Disease-associated variants are often enriched in tissue-specific enhancer-dense regions in disease-relevant cell types [54]. These regions assume a complex chromatin structure through DNA looping, regulating the expression of multiple genes. Assessed by HiC [34] and promoter-HiC [11] (Figure 2A and S3A), the enhancer-rich domain at the *SLC30A8* locus forms a unique 3D structure that has close contacts with multiple genes at both 5’ and 3’ ends. Although we did not detect significant associations between variants and nearby gene expression in cASE or eQTL analysis, we found that forced changes in enhancer structure imposed by genomic editing in EndoC-βH3 cells affected the expression of nearby genes.

We focused on two variant-bearing regulatory regions (R1 and R2 enhancer elements, Figure 2A) located in a single super-enhancer element embedded in the large enhancer-rich domain. These two regions are likely to exert the effects of T2D risk variants on *SLC30A8* and nearby gene expression, and contain β-cell specific transcription factor binding from FOXA2 and NKX2.2 (for R1) or PDX1 (for R2) (Figure S3B) (http://pasqualilab.upf.edu/app/isletregulome) [32]. Although neither region displayed transcriptional activity *per se* (Figure S5A-S5C and S6A-S6C), deletion of either R1 and R2 led to reduced expression of *SLC30A8* and other nearby genes examined (Figure 4B and 5B). These data demonstrate that these regions are relevant to activate gene expression by the super-enhancer domain and in turn impact the expression of multiple genes. It is possible that these regions play other role(s) than classical transcriptional regulation. Indeed, the wider variant region including R1 and R2 is spatially associated with CTCF-binding sites (CBSs) at both the 5’ and 3’ ends of the super-enhancer (Figure 2B). Thus, they may be involved in maintaining the chromatin structure of the enhancer-rich region. We have previously reported such a role for variants in the *STARD10* locus [17].

### Enhancer-bearing variants affect transcriptional activity and alter the expression of SLC30A8 and other genes

We focused here on three non-coding T2D GWAS variants and assessed their role in β cells. Variants located within a single super-enhancer are expected to alter transcriptional activity and, in turn, influence the levels of downstream transcript(s). The T2D protective alleles of rs3802177 and rs35859536 exhibited lower transcriptional activities in promoter-luciferase assays (Figure 6A and 6B) which is consistent with the cASE results. Furthermore, disruption of two variants, rs13266634 and rs35859538, led to significant changes on the expression levels of *SLC30A8* and nearby genes. These data suggest that these variants could affect the overall regulatory potential of the super-enhancer region. As listed in Table 4, the variant sites are likely bound by different transcription factor(s) between T2D risk and protective variants, potentially changing the nearby transcriptional landscape. We note that, of these variants, which are in LD to each other, rs13266634 hosts the majority of the posterior probability in the credible set (Table 3) [3], and as such it is unclear to what extent the neighbouring variants – and their independent effects on gene expression – contribute to T2D risk. In any case, future work will be needed to determine whether differential transcription factor binding in this region impacts local chromatin structure and enhancer activity.

### Genes regulated by the super-enhancer are critical for β-cell survival

Previous studies [12], and the present work, are consistent with the view that variants in *SLC30A8* influence transcriptional activity. The observation that protective variants are associated with lower *SLC30A8* mRNA levels suggests that this genetic regulatory layer could also influence the transcriptional activity of nearby genes. These include *RAD21*, *MED30* and *UTP23*, which are involved in chromatin organization [36], transcription [38] and translation [37], respectively. Correspondingly, we show that inactivation of these three genes exerts deleterious effects on β cell survival. However, the effects of subtle or more transient downregulation, which might conceivably exert positive effects on cell survival or function, were not explored.

Inactivation of *SLC30A8* had no evident impact on L-cell survival under the conditions used here, suggesting that effects of altered *SLC30A8* levels are tightly linked to acute effects on insulin secretion. Nevertheless, we note that, in a recent study [28], inactivation of *SLC30A8* in human stem cell-derived L-like cells (or the introduction of the loss-of-function R138X mutation) was protective against apoptosis under conditions of limiting Zn^2+^. Likewise, ZnT8 inactivation in mice protected islets from hypoxia and treatment with cytotoxic cytokines [55]. These earlier reports indicate that *SLC30A8* may influence cell viability under stressed conditions which pertain in diabetes.

### Targeting enhancers as a therapeutic approach to improve β cell function

Super-enhancers are characterized by strong transcription factor, co-factor, and enhancer-associated epigenetic modifications occupancy, and largely impact on the expression of key cell-identity genes [56]. We show that this high-density enhancer region not only drives β cell specific expression of the *SLC30A8*, but also regulates multiple nearby gene expression through complex chromatin looping. Furthermore, deletion of variant-bearing enhancers which are located in a single islet-selective super enhancer at the *SLC30A8* locus reduces the expression of all genes examined and improves insulin secretion. These data demonstrate that the super enhancer is a key regulator of β cell transcription and that targeted manipulation of this enhancer activity may conceivably alter β cell function in a therapeutic setting.

Super enhancers are enriched in genomic regions that harbour disease-associated genetic variants and are therefore relevant for clinical diagnosis and the identification of therapeutic targets [5, 32]. Examining both genetic data (cASE analysis) data and our human beta cell model, we show that T2D risk alleles within the *SLC30A8* enhancer cluster are associated with elevated transcriptional activity. Similar findings were also identified in other GWAS-identified genetic loci. For example, the risk allele of rs7163757, located in an active enhancer region, also increases the expression of nearby genes *C2CD4A* and *C2CD4B* [45].

JQ1 is the lead and most widely studied BET protein inhibitor. JQ1 competes for the acetyl-lysine binding pocket of the bromodomain, displacing the BET protein from chromatin binding, which alters the transcriptional activity of the target gene(s). Consistent with previous work [49], we found that JQ1 improves L cell function by increasing insulin content and secretion in EndoC-βH3 cells. Clarification of whether these effects are mediated by a reduction of the transcriptional activity of the enhancer cluster at the *SLC30A8* locus, or through changes in the expression of many other genes likely to be affected by this drug will, however, require future studies.

## Conclusions

We examined the role in pancreatic β cells of a previously identified islet enhancer cluster at the *SLC30A8* locus. We demonstrate that this enhancer domain regulates the expression of *SLC30A8,* and that T2D risk variants at this locus increase *SLC30A8* levels. The expression of nearby genes involved in fundamental cellular processes is also influenced by perturbations of this region *in vitro*. Future studies will be needed to determine the role of the latter genes in the effects of T2D variants and to assess the relative importance of changes in *SLC30A8* expression versus intrinsic Zn^2+^ transporter activity of the ZnT8 protein.

## Materials and Methods

### Expression quantitative trait loci (eQTL) analysis

Expression quantitative trait *loci* analyses were performed using two previously-published datasets [52, 57]. For Khamis et al., 2019, a total of 100 organ donors and 103 living donors of European descent were included in the study, with genotyping data (2.5M Omniarray Beadchip) from blood. We utilised the Genome-studio software to call the genotypes and QC thresholds for SNPs were as follows: a Hardy-Weinberg equilibrium > 0.001, a minor allele frequency > 0.05 and a call rate > 0.95. Imputation was performed using Impute2 (v2.3.2) from the 1,000 genomes panel (phase 3). Following QC, > 8.7M SNPs remained for further analysis. Ethnicity was tested using principal component analysis from 1,000 genomes and confirmed the European descent of subjects, therefore, population structure was not adjusted for. For the organ donors, RNA was isolated using collagenase, and in living donors isolated using laser capture microdissection (LCM). RNA gene expression was determined using Affymetrix (Human Genome U133 Plus 2.0 Array). RNA expression data was normalised using the Robust Multichip Analysis (RMA) method utilising the package affy [53] and transcriptomic batch effect was corrected for using the *combat* approach using the sva R package [58]. Genotyping and RNA data was integrated using the fastQTL software to identify eQTLs, adjusted for sex and age. P-values were computed using a permutation pass, with the number set from 1,000 to 10,000. Multiple testing was accounted for using Benjamini-Hochberg method to correct p-values. We report the nominal p-values for the tested eQTLs. For Vinuela et al [52], we utilised data published from a total of 26 preparations of FAC-sorted pancreatic beta cells and genotyped using the 2.5M Omniarray Beadchip. The eQTL analysis was performed using fastQTL, with a cis-window of 1 Mb, and adjusting for sex, batch effect (for RNA sequencing), PCs (for genotyping) and laboratory origin of samples. P-values were determined based on the 1,000 permutations per gene.

### Combined allele-specific expression (cASE) of transcript

The analysis of combined allele-specific expression (cASE) in multiple samples was performed as described in [12]. In brief, in all samples that are heterozygous for genomic variants within transcribed regions (reporter variants), the number of RNA-seq reads containing either the reference or alternative alleles is quantified. Then, their binomial significances are used to calculate a weighted Z-score, which measures the significance of the allelic bias across all samples, using the RNA-seq read coverage of each sample as weight. Significance is assessed by comparing the obtained Z-score with a control distribution created using the same read counts, but randomly shuffled across heterozygous individuals 1000 times.

All genomic variants located in the same Topologically Associated Domain (TAD) of each reporter variant are assessed as candidate variants responsible for the cASE effect. To do so, all samples that have a heterozygous reporter are separated in two subgroups, based on whether they are heterozygous or homozygous for the candidate variant. The Z-score is then calculated for each subgroup, using the heterozygous reporter variant. If the Z-score of the heterozygous candidate subgroup is more significant than the reporter’s Z-score (which was calculated with all the samples), and if the Z-score of the homozygous candidate subgroup is non-significant, the variant is then proposed as a putative candidate for mediating the combined allele-specific expression effect.

### Cell culture

The human-derived β cell line EndoC-βH3 was grown on extracellular matrix (ECM; 1% v/v) and fibronectin (2 μg/ml)-coated plates or petri dishes in serum-free DMEM containing low glucose (1 g/L), 2% (w/v) albumin from bovine serum fraction V, 50 μM β-Mercaptoethanol, 10 mM nicotinamide, 5.5 μg/mL human transferrin, 6.7 ng/mL sodium selenite, penicillin (100 units/mL), and streptomycin (100 μg/mL) [59]. HEK293T cell was cultured in DMEM high glucose (4500 mg/L) medium supplemented with 10% fetal bovine serum, 6 mM L-glutamine, penicillin (100 μg/mL) and streptomycin (100 μg/mL).

Rat INS-1 (832/13) cells were cultured in RPMI 1640 media supplemented with 1 mM pyruvate, 10 mM HEPES, penicillin (100 μg/mL), 100 μg/ml streptomycin and 10% fetal bovine serum (FBS).

### Mapping of the SLC30A8 regulatory landscape

We examined epigenomic datasets and pcHi-C chromatin interactions in human pancreatic islets for the *SLC30A8* locus [11]. For the super-enhancer domain harbouring cASE signals and T2D risk variants (chr8:118183060-118260108), we assessed chromatin activity among a broad range of tissues and cell-types using chromatin states generated by EpiMap ([33]). We included in Figure S2A, EpiMap chromatin states that showed active enhancer activity (EnhA1, EnhA2, EnhG1, EnhG2, EnhWk) in the super-enhancer domain.

### Chromatin Immunoprecipitation

Chromatin Immunoprecipitation (CHIP) was carried out as described [17]. In brief, 1 × 10^6^ EndoC-βH3 cells were fixed with 1% (v/v) formaldehyde for 10 minutes and quenched with 1.25 mM glycine. Cells were then scraped and resuspended in cell lysis buffer (2% Triton-100, 1% SDS, 100 mM NaCl, 10 mM Tris-HCl, 1 mM EDTA). After 20 stokes of homogenization with a disposable pestle, cells were sonicated for 10 min. using Covaris^TM^ S220 to breakdown genomic DNA to 200-500 bp fragments. DNA/protein complexes were then precipitated with anti-CTCF antibody (EMD Millipore) or rabbit IgG conjugated with protein A and G beads. DNAs were then purified through Phenol/Chloroform extraction and Ethanol precipitation.

### CRISPR-Cas9-mediated genome editing

gRNA sequences were designed using the software provided by Broad Institute (https://portals.broadinstitute.org/gpp/public/analysis-tools/sgrna-design) [60]. To generate mutations in EndoC-βH3 cells, lentiviral constructs carrying a gRNA and humanized *S. pyogenes* Cas9 (*hsp*Cas9) were transfected into HEK293T cells together with packaging plasmids pMD2.G and psPAX2 using CaCl_2_ transfection protocol [61]. Scramble gRNAs were served as a SHAM control (Table S2). Next day, cells were treated with sodium butyrate (10 mM) for 8 hours before changing to fresh medium. The medium was collected twice in the next three days and subjected to ultracentrifugation (Optima XPN-100 Ultracentrifuge, Beckman Coulter) at 26,000 rpm for 2 hours at 4°C. The lentiviruses were collected from the bottom of the tube and titrated. Same number of viruses was used to transduce to the EndoC-βH3 cells (MOI = 6-8). Blasticidin (25 μg/ml) was added 72 hour after infection to select lentivirus-infected cells.

For deletion of genomic regions, two lentiviral vectors were used. The first one (plenti-RIP-Cas9-BSD) carried a gRNA, Cas9 and Blasticidin resistant genes) and anther vector carried second gRNA with Hygromycin resistant gene and GFP gene cassette). Both plasmids were co-transfected into HEK293T cells with packaging plasmids. After infection, cells were monitored by GFP positivity and selected by both Blasticidin and hygromycin at final concentration of 25 μg/ml and 200 μg/ml, respectively. All gRNA sequences in this study are listed in Table S2.

To measure deletion efficiency after CRISPR/Cas9 mediated genome editing, SYBR Green qPCR was deployed to detect wild-type allele using primers 1 and 2 from genomic DNA extracted from control and DNA deleted/mutated cells. *CXCL12* was served as an internal DNA copy number control (Key resources table). The efficiency was calculated as: [1-2ΔΔCt(del-CXCL12)/2ΔΔCt(WT-CXCL12)] x 100%. In addition, the relative values of DNA deletion or inversion was also measured using primer set 1+4 or 1+3 respectively. The primer sets were listed in Table S4.

### PCR and RT-qPCR

Fusion high fidelity Taq polymerase (Thermo Fisher Scientific) was used in all routine PCR reactions to avoid PCR errors. A typical PCR reaction was set as follow: 98°C for 30 s, then with 35 cycles at 98°C 10 s, 60°C 10 s and 72°C 15 s. The primer sets for genomic DNA amplification are listed in Table S3.

Total RNA from EndoC-βH3 cells was obtained using TRIzol reagent (Invitrogen). Total RNAs (2 μg) were then reverse-transcribed into first strand cDNAs using High-Capacity cDNA Reverse Transcription Kit (Thermo Fisher Scientific) according to the manufacturer’s instructions. Real-time PCR was performed on a 7500 Fast Real-Time PCR System using the Fast SYBR^TM^ Green master mix or Fast Taqman^TM^ master mix. The Taqman gene expression primer/probe sets (Thermos Fisher) are listed in Table S1. The SYBR^TM^ Green PCR primer sets are listed in Table S4. The experiment was performed in duplicate and repeated three times.

### Molecular cloning

Regulatory regions (R1 and R2), identified by integration of previously published human islet ATAC-seq and H3K27ac ChIP-seq datasets [11], were PCR-amplified from genomic DNA extracted from EndoC-βH3 cells with primer sets (Table S5) designed by Primer3-based software and cloned into pGL4.23[*luc*/minP] vector (Promega) between KpnI and XhoI restriction enzyme sites. Plasmid DNA was extracted using mini-prep plasmid extraction kit and/or Maxi-prep plasmid extraction kit (QIAGEN). The constructs were further confirmed by the Sanger sequencing.

### Transfection and luciferase assay

EndoC-βH3 cells were seeded at a density of 50,000 per well in 96-well plates. After 48 hours, 0.1 μg of luciferase constructs containing putative regulatory sequences were co-transfected with 1 ng of pRL-Renilla construct as internal control into EndoC-βH3 cells, using Lipofectamine 2000, according to manufacturer’s instruction. pGL4.23 empty vector was served as a control. 48 h later, transfected cells were washed once with PBS and lysed directly in passive cell lysis buffer (Promega). Cells were incubated on a rotating platform at room temperature for 10 min. to ensure complete lysis of cells, and then spun at 10,000 rpm for 10 min to remove cell debris. Supernatant was transferred into a fresh tube and used to measure luciferase activity with Dual-Luciferase Reporter Assay kit (Promega) on a Lumat LB9507 luminometer (Berthold Technologies). Firefly luciferase measurements were normalized to *Renilla* luciferase.

### Insulin secretion

EndoC-βH3 cells were seeded onto ECM/Fibronectin-coated 96-well plates at 7.0 xL10^4^ cells per well. Two days after seeding, cells were incubated overnight in a glucose starving medium (glucose-free DMEM supplemented with 2% Albumin from bovine serum fraction V, 50 μl 2-mercaptoethanol, 10 mM nicotinamide, 5.5 μg/ml transferrin, 6.7 ng/ml sodium selenite, 100 units/ml, penicillin, 100 μg/ml streptomycin and 2.8 mM glucose). The next morning cells were incubated for 1 h in Krebs-Ringer solution [0.2% BSA, 25% solution 1 (460 mM NaCl), 25% solution II (96 mM NaHCO3, 20 mM KCl and 4 mM MgCl2), 25% solution III (4 mM CaCl2), 10 mM HEPES] supplemented with 0.5 mM glucose. EndoC-βH3 cells were then incubated in the presence of low (0.5 mM) or high glucose (15 mM) with or without 0.5 mM IBMX. After incubation for 1 h, the supernatant was collected, placed onto ice and centrifuged for 5 min. at 3,000 rpm at 4°C. The supernatant was then transferred into a fresh tube. Cells were lysed in 50 μL cell lysis solution (TETG: 20 mM Tris pH 8.0, 1% Triton X-100, 10% glycerol, 137 mM NaCl, 2 mM EGTA). The lysate was then removed to a fresh tube and centrifuged at 3,000 rpm for 5 min at 4°C. Insulin content was measured using an insulin ultra-sensitive assay kit (Cisbio). Secreted insulin was normalized as percentage of total insulin content. Fold increase in glucose- or other stimuli-stimulated insulin secretion is expressed as a ratio in comparison with secretion at basal level (0.5 mM glucose). Insulin secretion assays were performed in duplicate with insulin measurement in duplicate as well.

### Transcription factor binding motif analysis

TF binding profile on genetic variants was carried out using JASPAR CORE program (http://jaspar.genereg.net) [43, 62]. The threshold of relative profile score was set up at 80%. The scores of potential transcription factors were compared between risk and protective variants and listed in Table 4.

### Cell Growth curve

1 x 10^5^ EndoC-βh3 cells were plated in 12 well plate and infected with lentiviruses carrying a gRNA targeting a specific gene and Cas9. Three days later, cells were treated with blasticidin (25 μg/ml) and counted as day 0. To quantify live cell numbers for each sample following the time course, crystal violet staining was applied. [63] In brief, cells were stained with crystal violet solution (0.5% crystal violet and 20% methanol solution for 30 min, rinsed extensively with H_2_O, and dried. Cell-associated dye was extracted with 1.0 ml 10% acetic acid. Aliquots were diluted 1:4 with H_2_O, transferred to 96-well microtiter plates, and the optical density at 590 nm was determined. Values were normalized to the optical density at day 0. Within an experiment, each point was determined in triplicate.

### JQ1 treatment

7 x 10^4^ INS1 (832/13) or EndoC-βH3 cells were plated in 96 well plate and treated with JQ1 at various concentrations (from 25 nM to 200 nM) for 3 or 4 days [49]. At the end of the treatment, cells were lysed in lysis buffer (TETG: 20 mM Tris pH 8.0, 1% Triton X-100, 10% glycerol, 137 mM NaCl, 2 mM EGTA) and insulin content was measured by insulin ultra-sensitive assay kit (Cisbio). Within an experiment, each point was determined in triplicate and the experiment was repeated at least three times. For RNA extraction, cells were seeded at 5 x 10^5^ (INS1) or 1 x 10^6^ (EndoC-βH3) respectively and treated with various JQ1 concentrations for either 3 or 4 days. Cells were lysed in 1 ml Trizol and total RNAs were extracted according to manufacturer’s instructions.

### Statistical analysis

Data are expressed as means ± SEM. Significance was tested by Student’s two-tailed t test (paired or none-paired, Mann-Whitney test for non-parametric data, and one- or two-way ANOVA with SIDAK multiple comparison test, as appropriate, using Graphpad Prism 9.0 software. p < 0.05 was considered significant. The statistical details can be found in the figure legends (indicated as *n* number).

## Supporting information

all supplemnentary figures

graphical abstract

## Acknowledgements

G.A.R. was supported by a Wellcome Trust Investigator award (212625/Z/18/Z); UKRI-Medical Research Council (MRC) Programme grant (MR/R022259/1), an NIH-NIDDK project grant (R01DK135268) a CIHR-JDRF Team grant (CIHR-IRSC TDP-186358 and JDRF 4-SRA-2023-1182-S-N) CRCHUM start-up funds, and an Innovation Canada John R. Evans Leader Award (CFI 42649). We thank Dr Jorge Ferrer (CRG, Barcelona) for helpful discussion and advice.

## Author contributions

M. Hu and G.A. Rutter conceived and designed the research; M. Hu, I. Kim, W. Peng and O. Sun performed the research and acquired the data. I. Morán and S. Bonas-Guarch performed cASE and epigenomic analysis, respectively. A. Bonnefond, A. Khamis, P. Froguel and S. Bonas-Guarch performed eQTL analysis. M. Hu, S. Bonas-Guarch and G.A. Rutter wrote the manuscript with input from all of the authors.

## Data availability

*Functional studies:* Source data for functional studies are available upon request. Human genetic data is available as indicated in the source references from which they are taken.

## Conflict of Interest

GAR has received grant funding from, and is a consultant for, Sun Pharmaceuticals Inc.

## Notes

### Summary of Updates

We have refined the definitions of a super-enhancer where required, and added new data describing the effects of using a transcriptional regulator, JQ1. New authors have been added

## REFERENCES

1. Fuchsberger C, Flannick J, Teslovich TM, et al (2016) The genetic architecture of type 2 diabetes. Nature 536(7614):41–47. 10.1038/nature18642

2. Mahajan A, Taliun D, Thurner M, et al (2018) Fine-mapping type 2 diabetes loci to single-variant resolution using high-density imputation and islet-specific epigenome maps. Nat Genet 50(11):1505–1513. 10.1038/s41588-018-0241-6

3. Mahajan A, Spracklen CN, Zhang W, et al (2022) Multi-ancestry genetic study of type 2 diabetes highlights the power of diverse populations for discovery and translation. Nat Genet 54(5):560–572. 10.1038/s41588-022-01058-3

4. Yj K, S M, My H, et al (2022) The contribution of common and rare genetic variants to variation in metabolic traits in 288,137 East Asians. Nature communications 13(1). 10.1038/s41467-022-34163-2

5. Parker SCJ, Stitzel ML, Taylor DL, et al (2013) Chromatin stretch enhancer states drive cell-specific gene regulation and harbor human disease risk variants. Proc Natl Acad Sci USA 110(44):17921–17926. 10.1073/pnas.1317023110

6. Pasquali L, Gaulton KJ, Rodríguez-Seguí SA, et al (2014) Pancreatic islet enhancer clusters enriched in type 2 diabetes risk-associated variants. Nat Genet 46(2):136–143. 10.1038/ng.2870

7. Viñuela A, Varshney A, van de Bunt M, et al (2020) Genetic variant effects on gene expression in human pancreatic islets and their implications for T2D. Nat Commun 11(1):4912. 10.1038/s41467-020-18581-8

8. Greenwald WW, Chiou J, Yan J, et al (2019) Pancreatic islet chromatin accessibility and conformation reveals distal enhancer networks of type 2 diabetes risk. Nat Commun 10:2078. 10.1038/s41467-019-09975-4

9. Miguel-Escalada I, Bonàs-Guarch S, Cebola I, et al (2019) Human pancreatic islet three-dimensional chromatin architecture provides insights into the genetics of type 2 diabetes. Nat Genet 51(7):1137–1148. 10.1038/s41588-019-0457-0

10. Mahajan A, Taliun D, Thurner M, et al (2018) Fine-mapping type 2 diabetes loci to single-variant resolution using high-density imputation and islet-specific epigenome maps. Nat Genet 50(11):1505–1513. 10.1038/s41588-018-0241-6

11. Miguel-Escalada I, Bonàs-Guarch S, Cebola I, et al (2019) Human pancreatic islet three-dimensional chromatin architecture provides insights into the genetics of type 2 diabetes. Nat Genet 51(7):1137–1148. 10.1038/s41588-019-0457-0

12. Alonso L, Piron A, Morán I, et al (2021) TIGER: The gene expression regulatory variation landscape of human pancreatic islets. Cell Rep 37(2):109807. 10.1016/j.celrep.2021.109807

13. Atla G, Bonàs-Guarch S, Cuenca-Ardura M, et al (2022) Genetic regulation of RNA splicing in human pancreatic islets. Genome Biol 23(1):196. 10.1186/s13059-022-02757-0

14. da Silva Xavier G, Loder MK, McDonald A, et al (2009) TCF7L2 regulates late events in insulin secretion from pancreatic islet beta-cells. Diabetes 58(4):894–905. 10.2337/db08-1187

15. Hodson DJ, Mitchell RK, Marselli L, et al (2014) ADCY5 couples glucose to insulin secretion in human islets. Diabetes 63(9):3009–3021. 10.2337/db13-1607

16. Carrat GR, Hu M, Nguyen-Tu M-S, et al (2017) Decreased STARD10 Expression Is Associated with Defective Insulin Secretion in Humans and Mice. Am J Hum Genet 100(2):238–256. 10.1016/j.ajhg.2017.01.011

17. Hu M, Cebola I, Carrat G, et al (2021) Chromatin 3D interaction analysis of the STARD10 locus unveils FCHSD2 as a regulator of insulin secretion. Cell Reports 34(5):108703. 10.1016/j.celrep.2021.108703

18. Sladek R, Rocheleau G, Rung J, et al (2007) A genome-wide association study identifies novel risk loci for type 2 diabetes. Nature 445(7130):881–885. 10.1038/nature05616

19. Davidson HW, Wenzlau JM, O’Brien RM (2014) ZINC TRANSPORTER 8 (ZNT8) AND BETA CELL FUNCTION. Trends Endocrinol Metab 25(8):415–424. 10.1016/j.tem.2014.03.008

20. Rutter GA, Chimienti F (2015) SLC30A8 mutations in type 2 diabetes. Diabetologia 58(1):31–36. 10.1007/s00125-014-3405-7

21. Nicolson TJ, Bellomo EA, Wijesekara N, et al (2009) Insulin storage and glucose homeostasis in mice null for the granule zinc transporter ZnT8 and studies of the type 2 diabetes-associated variants. Diabetes 58(9):2070–2083. 10.2337/db09-0551

22. Weijers RN (2010) Three-dimensional structure of beta-cell-specific zinc transporter, ZnT-8, predicted from the type 2 diabetes-associated gene variant SLC30A8 R325W. Diabetol Metab Syndr 2(1):33. 10.1186/1758-5996-2-33

23. Merriman C, Huang Q, Rutter GA, Fu D (2016) Lipid-tuned Zinc Transport Activity of Human ZnT8 Protein Correlates with Risk for Type-2 Diabetes. J Biol Chem 291(53):26950–26957. 10.1074/jbc.M116.764605

24. Lemaire K, Ravier MA, Schraenen A, et al (2009) Insulin crystallization depends on zinc transporter ZnT8 expression, but is not required for normal glucose homeostasis in mice. Proc Natl Acad Sci USA 106(35):14872–14877. 10.1073/pnas.0906587106

25. Pound LD, Sarkar SA, Benninger RKP, et al (2009) Deletion of the mouse Slc30a8 gene encoding zinc transporter-8 results in impaired insulin secretion. Biochem J 421(3):371–376. 10.1042/BJ20090530

26. Flannick J, Thorleifsson G, Beer NL, et al (2014) Loss-of-function mutations in SLC30A8 protect against type 2 diabetes. Nat Genet 46(4):357–363. 10.1038/ng.2915

27. Dwivedi OP, Lehtovirta M, Hastoy B, et al (2019) Loss of ZnT8 function protects against diabetes by enhanced insulin secretion. Nat Genet 51(11):1596–1606. 10.1038/s41588-019-0513-9

28. Sui L, Du Q, Romer A, et al (2023) ZnT8 Loss of Function Mutation Increases Resistance of Human Embryonic Stem Cell-Derived Beta Cells to Apoptosis in Low Zinc Condition. Cells 12(6):903. 10.3390/cells12060903

29. Pickrell JK (2014) Joint analysis of functional genomic data and genome-wide association studies of 18 human traits. Am J Hum Genet 94(4):559–573. 10.1016/j.ajhg.2014.03.004

30. Mahajan A, Wessel J, Willems SM, et al (2018) Refining the accuracy of validated target identification through coding variant fine-mapping in type 2 diabetes. Nat Genet 50(4):559–571. 10.1038/s41588-018-0084-1

31. Costanzo MC, Grotthuss M von, Massung J, et al (2023) The Type 2 Diabetes Knowledge Portal: An open access genetic resource dedicated to type 2 diabetes and related traits. Cell Metabolism 35(4):695–710.e6. 10.1016/j.cmet.2023.03.001

32. Pasquali L, Gaulton KJ, Rodríguez-Seguí SA, et al (2014) Pancreatic islet enhancer clusters enriched in type 2 diabetes risk-associated variants. Nat Genet 46(2):136–143. 10.1038/ng.2870

33. Boix CA, James BT, Park YP, Meuleman W, Kellis M (2021) Regulatory genomic circuitry of human disease loci by integrative epigenomics. Nature 590(7845):300–307. 10.1038/s41586-020-03145-z

34. Greenwald WW, Chiou J, Yan J, et al (2019) Pancreatic islet chromatin accessibility and conformation reveals distal enhancer networks of type 2 diabetes risk. Nature Communications 10(1):2078. 10.1038/s41467-019-09975-4

35. Lawlor N, Stitzel ML (2019) (Epi)genomic heterogeneity of pancreatic islet function and failure in type 2 diabetes. Mol Metab 27S:S15–S24. 10.1016/j.molmet.2019.06.002

36. Cheng H, Zhang N, Pati D (2020) Cohesin subunit RAD21: From biology to disease. Gene 758:144966. 10.1016/j.gene.2020.144966

37. Wells GR, Weichmann F, Sloan KE, Colvin D, Watkins NJ, Schneider C (2017) The ribosome biogenesis factor yUtp23/hUTP23 coordinates key interactions in the yeast and human pre-40S particle and hUTP23 contains an essential PIN domain. Nucleic Acids Res 45(8):4796–4809. 10.1093/nar/gkw1344

38. Shukla A, Srivastava S, Darokar J, Kulshreshtha R (2020) HIF1α and p53 Regulated MED30, a Mediator Complex Subunit, is Involved in Regulation of Glioblastoma Pathogenesis and Temozolomide Resistance. Cell Mol Neurobiol. 10.1007/s10571-020-00920-4

39. Bernelot Moens SJ, Mooij HL, Hassing H. C, et al (2014) Carriers of Loss-of-Function Mutations in EXT Display Impaired Pancreatic Beta-Cell Reserve Due to Smaller Pancreas Volume. PLoS One 9(12). 10.1371/journal.pone.0115662

40. Xiao S, Huang Q, Ren H, Yang M (2021) The mechanism and function of super enhancer RNA. Genesis 59(5–6):e23422. 10.1002/dvg.23422

41. Merkenschlager M, Odom DT (2013) CTCF and Cohesin: Linking Gene Regulatory Elements with Their Targets. Cell 152(6):1285–1297. 10.1016/j.cell.2013.02.029

42. Weth O, Renkawitz R (2011) CTCF function is modulated by neighboring DNA binding factors. Biochem Cell Biol 89(5):459–468. 10.1139/o11-033

43. Castro-Mondragon JA, Riudavets-Puig R, Rauluseviciute I, et al (2022) JASPAR 2022: the 9th release of the open-access database of transcription factor binding profiles. Nucleic Acids Research 50(D1):D165–D173. 10.1093/nar/gkab1113

44. Kulzer JR, Stitzel ML, Morken MA, et al (2014) A common functional regulatory variant at a type 2 diabetes locus upregulates ARAP1 expression in the pancreatic beta cell. Am J Hum Genet 94(2):186–197. 10.1016/j.ajhg.2013.12.011

45. Kycia I, Wolford BN, Huyghe JR, et al (2018) A Common Type 2 Diabetes Risk Variant Potentiates Activity of an Evolutionarily Conserved Islet Stretch Enhancer and Increases C2CD4A and C2CD4B Expression. Am J Hum Genet 102(4):620–635. 10.1016/j.ajhg.2018.02.020

46. Belkina AC, Denis GV (2012) BET domain co-regulators in obesity, inflammation and cancer. Nat Rev Cancer 12(7):465–477. 10.1038/nrc3256

47. Donati B, Lorenzini E, Ciarrocchi A (2018) BRD4 and Cancer: going beyond transcriptional regulation. Molecular Cancer 17(1):164. 10.1186/s12943-018-0915-9

48. Filippakopoulos P, Qi J, Picaud S, et al (2010) Selective inhibition of BET bromodomains. Nature 468(7327):1067–1073. 10.1038/nature09504

49. Deeney JT, Belkina AC, Shirihai OS, Corkey BE, Denis GV (2016) BET Bromodomain Proteins Brd2, Brd3 and Brd4 Selectively Regulate Metabolic Pathways in the Pancreatic β-Cell. PLoS One 11(3):e0151329. 10.1371/journal.pone.0151329

50. Rutter GA, Chabosseau P, Bellomo EA, et al (2016) Intracellular zinc in insulin secretion and action: a determinant of diabetes risk? Proc Nutr Soc 75(1):61–72. 10.1017/S0029665115003237

51. Kleiner S, Gomez D, Megra B, et al (2018) Mice harboring the human SLC30A8 R138X loss-of-function mutation have increased insulin secretory capacity. Proc Natl Acad Sci USA 115(32):E7642–E7649. 10.1073/pnas.1721418115

52. Viñuela A, Varshney A, van de Bunt M, et al (2020) Genetic variant effects on gene expression in human pancreatic islets and their implications for T2D. Nat Commun 11(1):4912. 10.1038/s41467-020-18581-8

53. Atla G, Bonàs-Guarch S, Cuenca-Ardura M, et al (2022) Genetic regulation of RNA splicing in human pancreatic islets. Genome Biol 23(1):196. 10.1186/s13059-022-02757-0

54. Hnisz D, Abraham BJ, Lee TI, et al (2013) Super-enhancers in the control of cell identity and disease. Cell 155(4):934–947. 10.1016/j.cell.2013.09.053

55. Karsai M, Zuellig RA, Lehmann R, et al (2022) Lack of ZnT8 protects pancreatic islets from hypoxia- and cytokine-induced cell death. J Endocrinol 253(1):1–11. 10.1530/JOE-21-0271

56. Whyte WA, Orlando DA, Hnisz D, et al (2013) Master transcription factors and mediator establish super-enhancers at key cell identity genes. Cell 153(2):307–319. 10.1016/j.cell.2013.03.035

57. Khamis A, Canouil M, Siddiq A, et al (2019) Laser capture microdissection of human pancreatic islets reveals novel eQTLs associated with type 2 diabetes. Mol Metab 24:98–107. 10.1016/j.molmet.2019.03.004

58. Leek JT, Johnson WE, Parker HS, Jaffe AE, Storey JD (2012) The sva package for removing batch effects and other unwanted variation in high-throughput experiments. Bioinformatics 28(6):882–883. 10.1093/bioinformatics/bts034

59. Benazra M, Lecomte M-J, Colace C, et al (2015) A human beta cell line with drug inducible excision of immortalizing transgenes. Mol Metab 4(12):916–925. 10.1016/j.molmet.2015.09.008

60. Doench JG, Fusi N, Sullender M, et al (2016) Optimized sgRNA design to maximize activity and minimize off-target effects of CRISPR-Cas9. Nat Biotechnol 34(2):184–191. 10.1038/nbt.3437

61. Graham FL, van der Eb AJ (1973) Transformation of rat cells by DNA of human adenovirus 5. Virology 54(2):536–539. 10.1016/0042-6822(73)90163-3

62. Khan A, Fornes O, Stigliani A, et al (2018) JASPAR 2018: update of the open-access database of transcription factor binding profiles and its web framework. Nucleic Acids Res 46(D1):D260–D266. 10.1093/nar/gkx1126

63. Serrano M, Lin AW, McCurrach ME, Beach D, Lowe SW (1997) Oncogenic ras provokes premature cell senescence associated with accumulation of p53 and p16INK4a. Cell 88(5):593–602. 10.1016/s0092-8674(00)81902-9

64. Guyonnet S, Rolland Y, Takeda C, et al (2021) The INSPIRE Bio-Resource Research Platform for Healthy Aging and Geroscience: Focus on the Human Translational Research Cohort (The INSPIRE-T Cohort). J Frailty Aging 10(2):110–120. 10.14283/jfa.2020.38

65. Lawlor N, Márquez EJ, Orchard P, et al (2019) Multiomic Profiling Identifies cis-Regulatory Networks Underlying Human Pancreatic β Cell Identity and Function. Cell Rep 26(3):788–801.e6. 10.1016/j.celrep.2018.12.083

