## Supplementary figure legends, supplementary tables for "Multiple genetic variants at the *SLC30A8* locus affect local super-enhancer activity and influence pancreatic β-cell survival and function"

Supplementary Information

Supplementary Figures

Figure S1 **GWAS identified genetic variants at the *SLC30A8* gene locus.** A. The GWAS variants (red dots) are located at the 3’ end of the SLC30A8 gene. B. Islet-specific transcriptional factor binding sites at the SLC30A8 locus. R1 and R2 (shown in red) are two active enhancer regions bearing 3 GWAS variants. R1 and R2 in red are two enhancer regions bearing GWAS variants.

Figure S2 **Epigenomic map at the *SLC30A8* locus in pancreatic islet and other cell types.** ChromHMM states were extracted from EpiMap (<http://compbio.mit.edu/epimap>).

Figure S3 **Detailed locations of R1 and R2 regions at the *SLC30A8* locus.** R1 region bearing rs13266634 and rs3802177 is overlap with the exon8 of the *SLC30A8* gene while R2 bearing rs35859536 is downstream of the *SLC30A8* transcript.

Figure S4 **Mutation of CTCF-binding sites (CBSs) in EndoC-bH3 cells.** A. Sanger sequencing of PCR product amplified from genomic DNA in wildtype and CBS-mut cells. Red box: PAM sequence; Red arrow: Cas9 cutting site. B. Comparison of CTCF binding affinity before and after CRISPR-Cas9 editing at CBSs. CTCF binding affinity was normalized to IgG control in wild-type and CBS-mut cells. C. Mutation efficiency of CRISPR-Cas9 genome editing in single CBS-mut cells. SYBR Green qPCR analysis was performed to detect wildtype DNA in control and CBS-mut cells using qPCR primers listed in Table S4. *n* = 2. D. Mutation efficiency of CRISPR-Cas9 genome editing in double CBS-mut cells. SYBR Green qPCR analysis was performed to detect wildtype DNA in control and double CBS-mut cells using qPCR primers listed in Table S4. *n* = 2.

Figure S5 **Role of R1 in β-cell function.** A. Diagram of R1 genomic DNA fragments in pGL4.23 luciferase vector. B and C. Promoter-luciferase assay in EndoC-βH3 cells. Data are mean ± SEM. *, *P* < 0.05; **, *P* < 0.01; ***, *P* < 0.005. *n* = 3. D. Sanger sequencing of PCR product amplified from R1-del1 cell. Red and blue bars represent the 5’ and 3’ end of DNA sequence flanking R1 region. It seems that the Cas9 cutting caused mutation(s) before ligation, which led to unclear DNA sequencing result. Primers used to amplify genomic DNA were listed in Table S3. E. Sanger sequencing of PCR product amplified from R1-del2 cell. Red and blue bars represent the 5’ and 3’ end of DNA sequence flanking R2 region. F. Diagram of qPCR strategy to determine the deletion efficiency. CXCL12 was served as an internal DNA copy number control. G. Deletion efficiency. The deletion efficiency rate (%) was determined by: 1. calculating the remaining wildtype allele (primer 1+4) of genomic DNA in R1-del cell and compared with wildtype allele in wild type control cell and 2. taking away the rate of DNA inversion (primers 1+3 or primers 2+3) in R1-del cell. *n* = 2.

Figure S6 **Role of R2 in β-cell function.** A. Diagram of R2 DNA fragments in pGL4.23 luciferase vector. B and C. Promoter-luciferase assay in EndoC-βH3 cells. Data are mean ± SEM. *, *P* < 0.05; **, *P* < 0.01; ***, *P* < 0.005. *n* = 3. D. Sanger sequencing of PCR product amplified from R2-del1 cell. E. Sanger sequencing of PCR product amplified from R2-del2 cell. F. Diagram of qPCR strategy to determine CRISPR-Cas9 mediated genomic DNA deletion efficiency. G. Deletion efficiency. *n* = 2.

Figure S7 **Analysis of individual variants in EndoC-H3 cells.** A. Sanger sequencing confirmation of luciferase vectors carrying either risk or protective variant. B. Sanger sequencing analysis of PCR product amplified from genomic DNA at variant site after CRISPR-Cas9 mediated genome editing. The editing resulted in mixed DNA sequences with no clear sequencing pattern at Cas9 cutting site in all three cell lines. C. Mutation efficiency of CRISPR-Cas9 mediated genome editing. The primer sets used for PCR amplification and DNA sequencing were listed in Table S3.

Figure S8 **Sanger sequencing of PCR product amplified from genomic DNA in wild-type and gene KO cells.** A. *RAD21*. B. *UTP23*. C. *MED30*. Note that the genomic DNAs were extracted from live cells. Since the knockout cells didn’t survive well, the sequencing results may not represent the KO effects. Nevertheless, some changes were observed at the Cas9 cutting site: *UTP23*-gRNA1: 1 bp (A) insertion and *UTP23*-gRNA2: 3 bp (CAA) deletion; *MED30*-gRNA1: 1 bp (G) insertion; *MED30*-gRNA2: 2 bp (GC) deletion; No significant change was observed in both *RAD21*-KO cells.

Figure S9 **Sanger sequencing of PCR product amplified from genomic DNA in wild-type and gene KO cells.** A. *EXT1*. B. *SLC30A8*. Sequencing changes were identified at the Cas9 cutting site as follows: EXT1-gRNA1: 1 bp (A) insertion; *EXT1*-gRNA2: mixed with no clear pattern; *SLC30A8*-gRNA1: 1 bp (C) deletion and SLC30A8-gRNA2: 1 bp (A) deletion. C. Statistical analysis of surviving cells presented in Figure 7A. 4 small areas on each plate were randomly selected and live cells were counted and compared with the Scrambled gRNA samples. Data are mean ± SEM. *, *P* < 0.05; **, *P* < 0.01; ***, *P* < 0.005; ****, *P* < 0.001.

Figure S10 **JQ1 increases insulin production and secretion in INS1 (832/13) cells**. A and B. Insulin content in JQ1 treated INS1 (832/13) cells. A. Total insulin content three days after JQ1 treatment in each well. Cells were seeded at 7 x 10^4^ cells in 96-well plate and incubated with JQ1 for three days. DMSO was served as a control. B. Fold change. Data are normalized to the insulin content in DMSO treated cells. *n* = 3. C. Taqman^TM^ RT-qPCR analysis of *Slc30A8* and nearby genes in JQ1 treated INS1 (832/13) cells. 5 x 10^5^ INS1 cells were seeded in 6-well plate and incubated with different concentrations of JQ1 for 3 days. Data are mean ± SEM. *, *P* < 0.05; **, *P* < 0.01; ***, *P* < 0.005; ****, *P* < 0.001.

**Supplementary Tables**

Table S1 Key Resources

| REAGENT or RESOURCE | SOURCE | IDENTIFIER |
| --- | --- | --- |
| Bacteria and Virus Strain | | |
| Stbl3 competent cells | Thermo Fisher Scientific | Cat# C737303 |
| 10-beta competent cells | New England Biolabs | Cat# C30191 |
| Chemicals, Peptides, and Recombinant Proteins | | |
| 3-Isobutyl-1-methylxanthine (IBMX) | Sigma-Aldrich | Cat# I5879 |
| Sodium Selenite | Sigma-Aldrich | Cat# S1382 |
| Nicotinamide | Sigma-Aldrich | Cat# 481907 |
| Paraformaldehyde | Agar Scientific | Cat# AGR1026 |
| TRIzol | Invitrogen | Cat# 15596018 |
| Albumin from Bovine Serum Fraction V | Roche Diagnostics | Cat# 10775835001 |
| Human Transferrin | Sigma-Aldrich | Cat# T8158 |
| ECM | Sigma-Aldrich | Cat# E1270 |
| Critical Commercial Assay | | |
| Dual-Luciferase Assay Reporter Assay System | Promega | Cat# E1910 |
| Lipofectamine 2000 | Thermo Fisher Scientific | Cat# 11668025 |
| Insulin Ultra-sensitive Assay Kit | Cisbio Bioassays | Cat# 62IN2PEH |
| High-Capacity cDNA Reverse Transcription Kit | Thermo Fisher Scientific | Cat# 4368814 |
| Taqman^TM^ Fast advanced master mix | Thermo Fisher Scientific | Cat# 4444553 |
| Fast SYBR^TM^ Green Master Mix | Thermo Fisher Scientific | Cat# 4385612 |
| Phusion High fidelity DNA polymerase | Thermo Fisher Scientific | Cat# F530 |
| JQ1 | Sigma-aldrich | SML1524-5MG |
| Experimental Model: cell line | | |
| EndoC-βH3 | Humancelldesign.com | N/A |
| Oligonucleotides | | |
| gRNAs | See Table S2 |  |
| Genomic DNA amplification | See Table S3 |  |
| SYBR^TM^ green qPCR for gene | See Table S4 |  |
| Luciferase assay | See Table S5 |  |
| Recombinant DNA | | |
| pMD2.G | Didier Trono (https://tronolab.epfl.ch) | Cat# 12259; RRID: Addgene_12259 |
| pSPAX2 | Didier Trono (https://tronolab.epfl.ch) | Cat# 12260; RRID: Addgene_12260 |
| pLenti-CRISPR-RIP-Cas9 | Hu, M. et al., Cell Reports (2021) | N/A |
| pGL4.23 | Promega | E8411 |
| pRL-Renilla | Promega | E2231 |
| Taqman^TM^ qPCR primer/probe | | |
| *SLC30A8* | Thermo Fisher Scientific | Cat# 4331182; Assay ID: Hs00545183_ml |
| *UTP23* | Thermo Fisher Scientific | Cat# 4331182; Assay ID: Hs00260536_ml |
| *RAD21* | Thermo Fisher Scientific | Cat# 4331182; Assay ID: Hs00366726_ml |
| *MED30* | Thermo Fisher Scientific | Cat# 4331182; Assay ID: HS00369804_ml |
| *EXT1* | Thermo Fisher Scientific | Cat# 4331182; Assay ID: Hs00609162_ml |
| *ACTB* | Thermo Fisher Scientific | Cat# 4331182; Assay ID: Hs01060665_gl |
| *EIF3H* | Thermo Fisher Scientific | Cat# 4331182; Assay ID: Hs00186779_m1 |
| *INS* | Thermo Fisher Scientific | Cat# 4331182; Assay ID: Hs00355773_m1 |
| *Actb* | Thermo Fisher Scientific | Cat# 8779616; Assay ID: Rn00667869_m1 |
| *Eif3h* | Thermo Fisher Scientific | Cat# 8777417; Assay ID: Rn01505699_g1 |
| *Utp23* | Thermo Fisher Scientific | Cat# 8777417; Assay ID: Rn01506218_m1 |
| *Rad21* | Thermo Fisher Scientific | Cat# 8777417; Assay ID: Rn01505656_m1 |
| *Med30* | Thermo Fisher Scientific | Cat# 8777417; Assay ID: Rn01200407_m1 |
| *Ext1* | Thermo Fisher Scientific | Cat# 8777417; Assay ID: Rn00468764_m1 |
| *Ins-1* | Thermo Fisher Scientific | Cat# 8777426; Assay ID: Rn02121433_g1 |
| *Ins-2* | Thermo Fisher Scientific | Cat# 8777426; Assay ID: Rn01774648_g1 |
| Software Thermo Fisher Scientific | | |
| GraphPad Prism 9 | GraphPad Software | RRID: SCR_002798 |

Table S2 Guide RNA (gRNA) sequences for CRISPR-Cas9 genome editing

| CBS1 | TACCACATTCGGCCTCAGGA |
| --- | --- |
| CBS2 | GGCGCCTTCTCGCCACAAGA |
| CBS3 | AGCGTGGGGGAACCACTAGA |
| CBS4 | TTTCCAGTGTGGCCACAAGA |
| CBS5 | TTCTTCCCAGCTGTCGGCAG |
| SCRAMBLED-gRNA1 | GAACTCAACCAGAGGGCCAA |
| SCRAMBLED-gRNA2 | GGGAGGTGGCTTTAGGTTTT |
| SLC30A8-gRNA1 | GCTCCAAGCCCACAGAAAAG |
| SLC30A8-gRNA2 | ATGAGTACGCCTATGCCAAG |
| RAD21-gRNA1 | CTAAATTACACTCGAACACA |
| RAD21-gRNA2 | TCTCAGTAAAAGAGGGCCTC |
| UTP23-gRNA1 | GCAGGATCTGGTACGGCTCG |
| UTP23-gRNA2 | CGCGGACTCCGAAGTTGTTG |
| MED30-gRNA1 | CCCCGACGCGGCCAACGGAG |
| MED30-gRNA2 | CACCCCTCCGTTGGCCGCGT |
| EXT1-gRNA1 | GCTTGCAGTTTAGGGCATCG |
| EXT1-gRNA2 | TGTGTTCTTCTCTCCGGCTG |
| R1-gRNA1 | GGGGCTTAGGTCTGACTTGA |
| R1-gRNA-2 | GCCTCTTCCTTCATGGTGAA |
| R1-gRNA3 | TTGTCTTTGACCTGCTGGGG |
| R2-gRNA3 | GGGGAGGATGCCTATAAGAT |
| R2-gRNA-2 | GTTTGGGGTAAACTGCAAAC |
| R2-gRNA-3 | TTGCTGGAGTGGTGTACATG |
| rs13266634-gRNA | AACCACTTGGCTGTCCCGGC |
| rs3802177-gRNA | AAAGGAAGAAATTCATGTCA |
| rs35859536-gRNA | TGGTAGATGGCTAGGTACGA |

Table S3 Primer set for PCR amplification of genomic DNA

| SLC30A8-F1 | AGCGGTTCCAAGTATGGGCAGG |
| --- | --- |
| SLC30A8-F2 | GCAGGGTGGGTCAGTGAGTGG |
| SLC30A8-R | GAAAGACTCACCCACGACCTCTGC |
| RAD21-F1 | CTCTAGCTCTTGGCCAGTAAACAGAC |
| RAD21-F2 | TGACTCTTGTCCCTTTTACTTTCAGCTC |
| RAD21-R | CTTGCACTCCAATGCCCCTACC |
| UTP23-F1 | GCTTCATTTCCGGGTGAAACTGGC |
| UTP23-F2 | GTGAAACTGGCATTGAGGGTACTGG |
| UTP23-R | CGCAATGAAGGTCTGTGAGCAACG |
| MED30-F1 | GCGGCCGCTGTTTTGAAATCG |
| MED30-F2 | GGGGTCTCTCAAGCTGGTTCC |
| MED30-R | CTTTCCCTCCCAGCTGCATCC |
| EXT1-F1 | GAAGTCTTTACAGGCGGGAAGATGG |
| EXT1-F2 | GGGAAGATGGCGGACTGGAGC |
| EXT1-R | ACTCCATGCGGCACTTCTTGC |
| R1-KO-F1 | AATTATGGTGGGGTCCCACACTG |
| R1-KO-F2 | GGCAGTGGGAGGGCATTTACTGC |
| R1-KO-R2 | CTGTTACTTCGGCTCCACTCAGG |
| R2-KO-F1 | TTTTCCTGTTTATTCCAGAACTTGGTAGG |
| R2-KO-F2 | CTCTCATTACTGAGATGTCAGTTTCTGC |
| R2-KO-R2 | CCTGGGGGAAGCTACAGTGG |
| CBS1-F1 | CACTTCCGCAGATGCATTCTGTGG |
| CBS1-F2 | CCGCAGATGCATTCTGTGGTAAGG |
| CBS1-R | AGGGAAATTTGGGGGCTAAAAGAGAGG |
| CBS2-F1 | TGCTTCTAGAGCAAATATTGAGCGTGG |
| CBS2-F2 | TTCGGGGAATCAGATCCTCCTGG |
| CBS2-R | CTCCCAGTTACACCGCTGAAAGG |
| CBS3-F1 | GCTGCTCTGTCATCTTGCAGACC |
| CBS3-F2 | CATCTTGCAGACCCACCCTGG |
| CBS3-R | CTTCTATGGCATTCCGCACCAG |
| CBS4-F1 | GCACTCACCTGTAGCCCCAGC |
| CBS4-F2 | GAGGGATCTGATCCCCCTACC |
| CBS4-R | GGGGATGTGAGGGAACACTCC |
| CBS5-F1 | AAGACAGTTGGCCAAAGTGATGAC |
| CBS5-F2 | CCTCCCACGCCCAAATCTGC |
| CBS5-R | CACACCCAACAAGACCCCAGC |

Table S4 SYBR^TM^ Green qPCR primer for lncRNA expression and genotyping

| β-ACTIN-F | CACCATTGGCAATGAGCGGTTC |
| --- | --- |
| β-ACTIN-R | AGGTCTTTGCGGATGTCCACGT |
| RAD21AS1-F | CTGTTTGTGGAGTCCACAACTGAGC |
| RAD21AS1-R | GTGTCCTCTTCAATTTCTTTCATCACTGG |
| RP11-654G14-F | AATATGCTCCCTGTGTTATTATTGCC |
| RP11-654G14-R | CCTAAACTCTCGACCAAGTCACC |
| rs13266634-F | GTCAGAGCAGTCGCCCATGC |
| rs13266634-R | CCACTTGGCTGTCCCGGCTGG |
| rs3802177-F | ACATGCTGCTATGCAGTTTCTGC |
| rs3802177-R | AAATGTGCATTGCACCATGACATG |
| rs35859536-F | GGTCGTGTGCCTGGTTCTAGACG |
| rs35859536-R | ATGCCAGATTGCCTCCATCGTAC |
| qR1-1 | CATTTGAAAACTGTGCTTTCTCAGACATCG |
| qR1-2 | CAAACGTGGCTTCCTCTGAGTGC |
| qR1-3 | GGAAGAAATTCATGTCATGGTGCAATGC |
| qR1-4 | TCCAATTGATTGATGGATCTCAGTGC |
| qR2-1 | CGGAAGTAGAGCCAAGAAAGATTAGAGG |
| qR2-2 | CTCAGAGAGAGAGGAGAGAATTGCTGG |
| qR2-3 | CAGCACTTCATTCTCTTTTCCTGCAG |
| qR2-4 | TGTGATGAGGCTGGTTCTTTGAGTTTGG |
| qCBS1-F | GGTGAAGTTAGAGAACACGCAGAGG |
| qCBS1-R | TAGATACCACATTCGGCCTCAGG |
| qCBS2-F | GTGGGGGAAATACTGCTGCAGG |
| qCBS2-R | GAAGGCGCCTTCTCGCCACAAG |
| qCBS3-F | CCCTCACCGCCGGTGTTCAG |
| qCBS3-R | GAGAGCGTGGGGGAACCACTAG |
| qCBS4-F | CTTAATTTCCAGTGTGGCCACAAG |
| qCBS4-R | CTACTGTGTGTCCATGCTCCCTGG |
| qCBS5-F | CTGGTCTGTGTCTCCAGCAAGC |
| qCBS5-R | GTCTTTCTTCCCAGCTGTCGGC |

Table S5 Primer set for DNA cloning into pGL4.23

| rs13266634-Luc-KpnI | ATGTCAGGTACCGTCAGAGCAGTCGCCCATGC |
| --- | --- |
| rs13266634-Luc-XhoI | AATCATCTCGAGCTGAATGGTGAGTGAGTGCATCG |
| rs3802177-Luc-KpnI | ATGTCAGGTACCGACTAGCTCAGTCACACCGTCAG |
| rs3802177-Luc-XhoI | AATCATCTCGAGCACATTCCTGCAATTGAACACAGC |
| rs35859536-Luc-KpnI | ATGTCAGGTACCGTCGTGTGCCTGGTTCTAGACG |
| rs35859536-Luc-XhoI | AATCATCTCGAGGATGACGATTTCACTCTTGCTCC |
| R1-Luc-KpnI-F1 | ATGTCAGGTACCGGGGCTTAGGTCTGACTTGAGG |
| R1-Luc-KpnI-F2 | ATGTCAGGTACCCAGAGATCCCTGGGAGCTGG |
| R2-Luc-KpnI-F1 | ATGTCAGGTACCCTGCTAGTGCTGGAATAGATCTCGG |
| R2-Luc-XhoI-R | AATCATCTCGAGTGTGATGAGGCTGGTTCTTTGAGTTTGG |

**Abbreviations**

T2D, Type 2 Diabetes

GWAS, Genome-wide association study

R, Regulatory region

SNP, Single-nucleotide polymorphism

CHIP-seq: Chromatin immunoprecipitation and DNA sequencing

ATAC-seq, Assay for Transposase-Accessible Chromatin using sequencing

3C, Chromosome Conformation Capture

4C, Circular Chromosome Conformation Capture

CTCF, CCCTC-bind factor

pcHi-C, Promoter Capture Hi-C

TAD, Topologically Associating Domain

SLC30A8, Solute Carrier Family 30 Member 8

RAD21, Double-strand-break repair protein rad21 homolog

UTP23, UTP23 Small Subunit Processome Component

MED30, Mediator of RNA Polymerase II Transcription Subunit 30

EXT1, Exostosin Glycosyltransferase 1

CRISPR, Clustered Regularly Interspaced Short Palindromic Repeats

Cas9, Endonuclease from *Streptococcus pyogenes*

GSIS, Glucose-stimulated insulin secretion
