## Supplementary figures and images for "Multiple genetic variants at the *SLC30A8* locus affect local super-enhancer activity and influence pancreatic β-cell survival and function"

### all supplemnentary figures

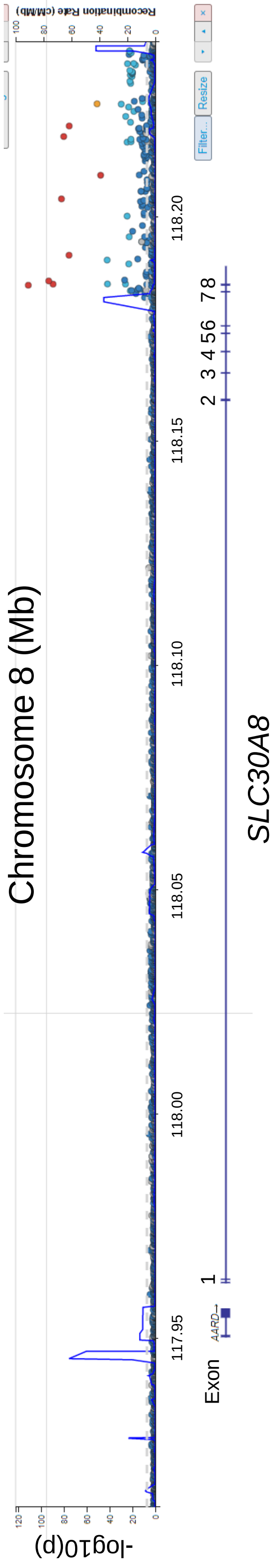

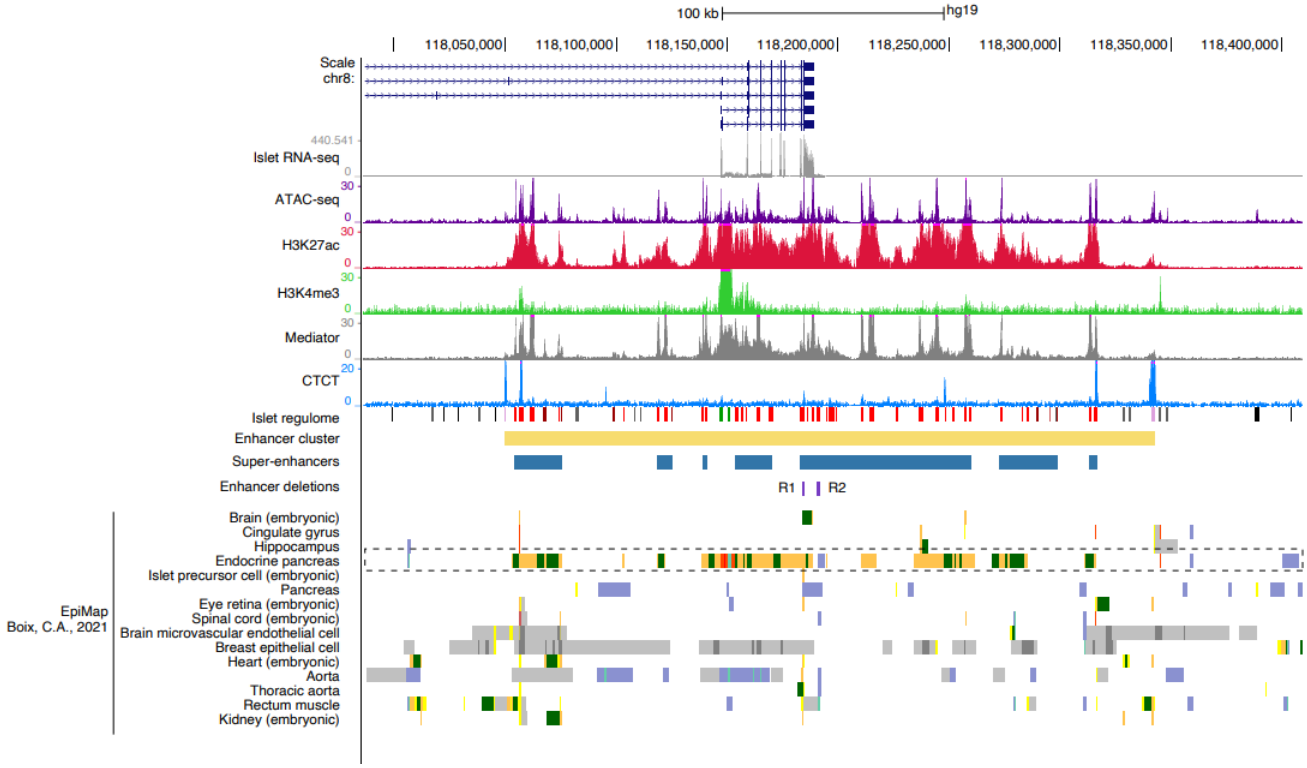

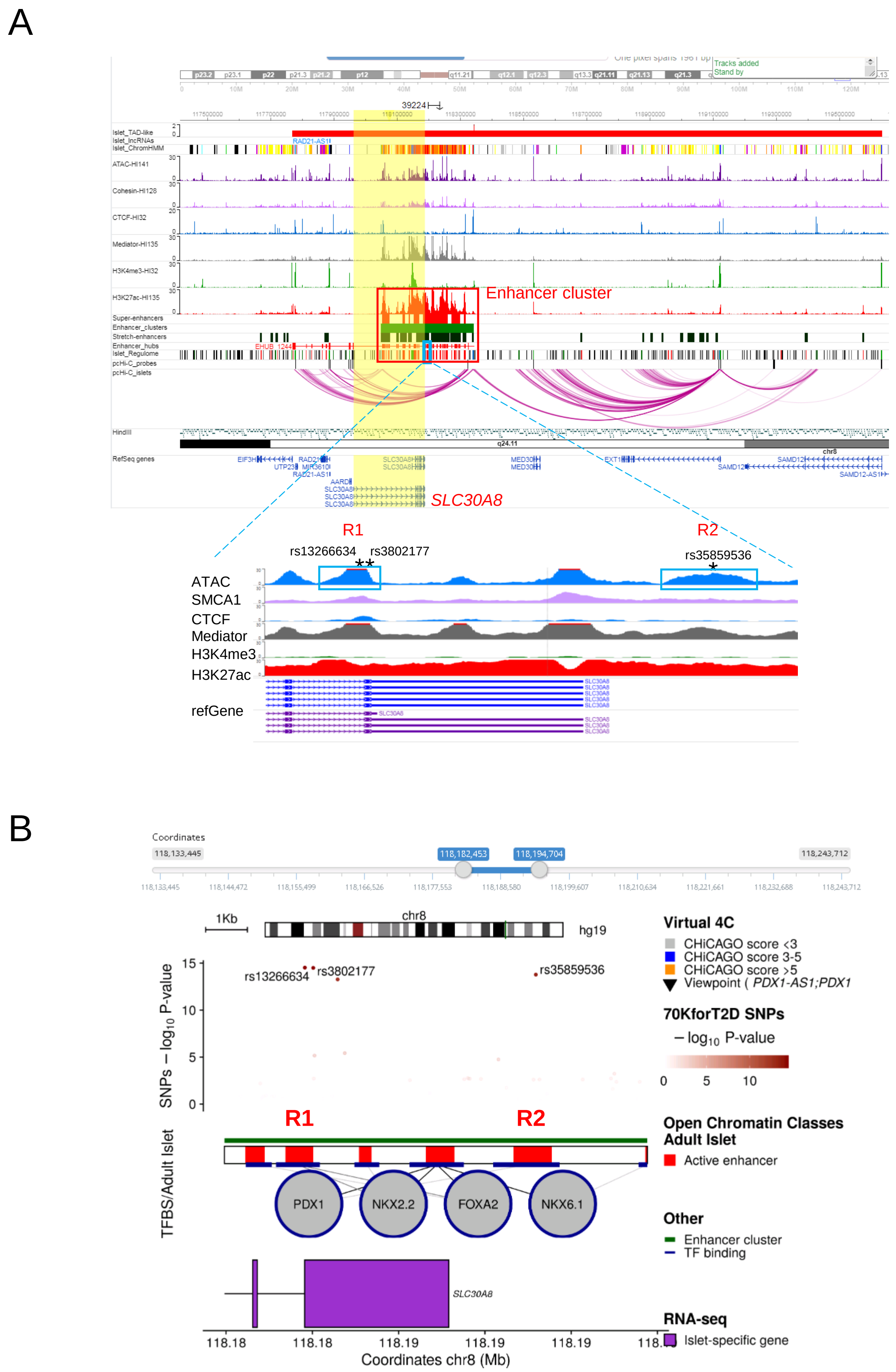

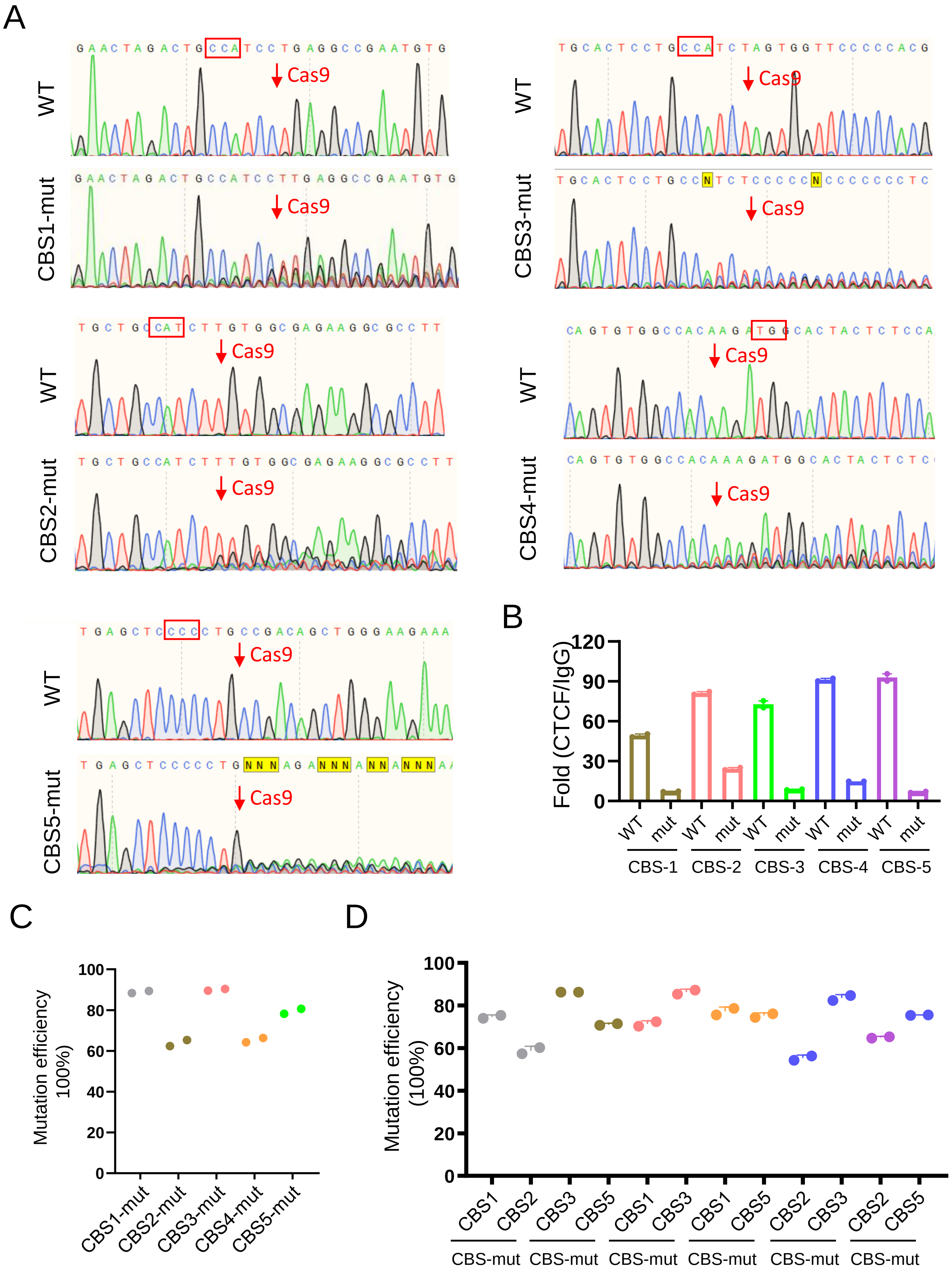

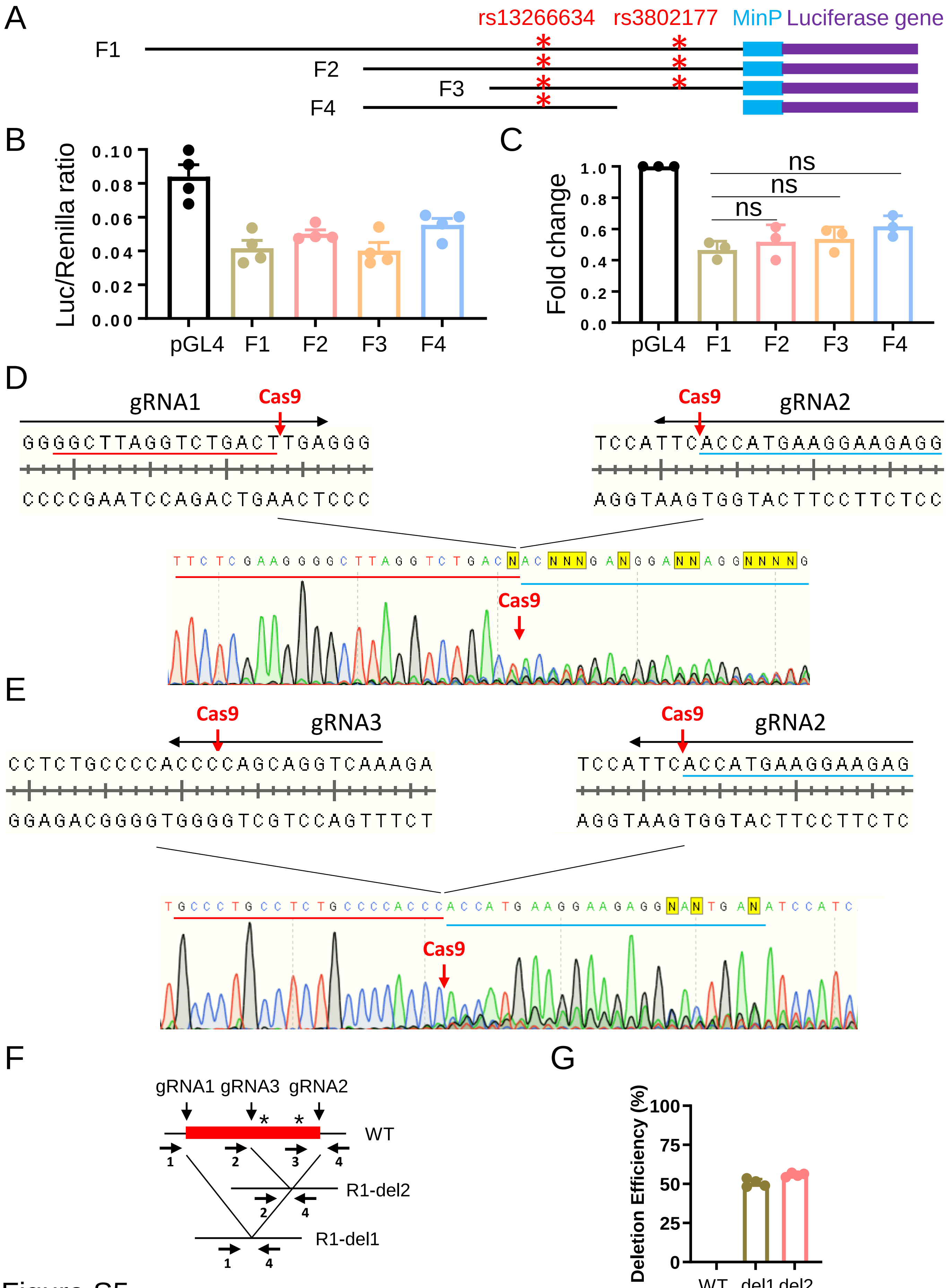

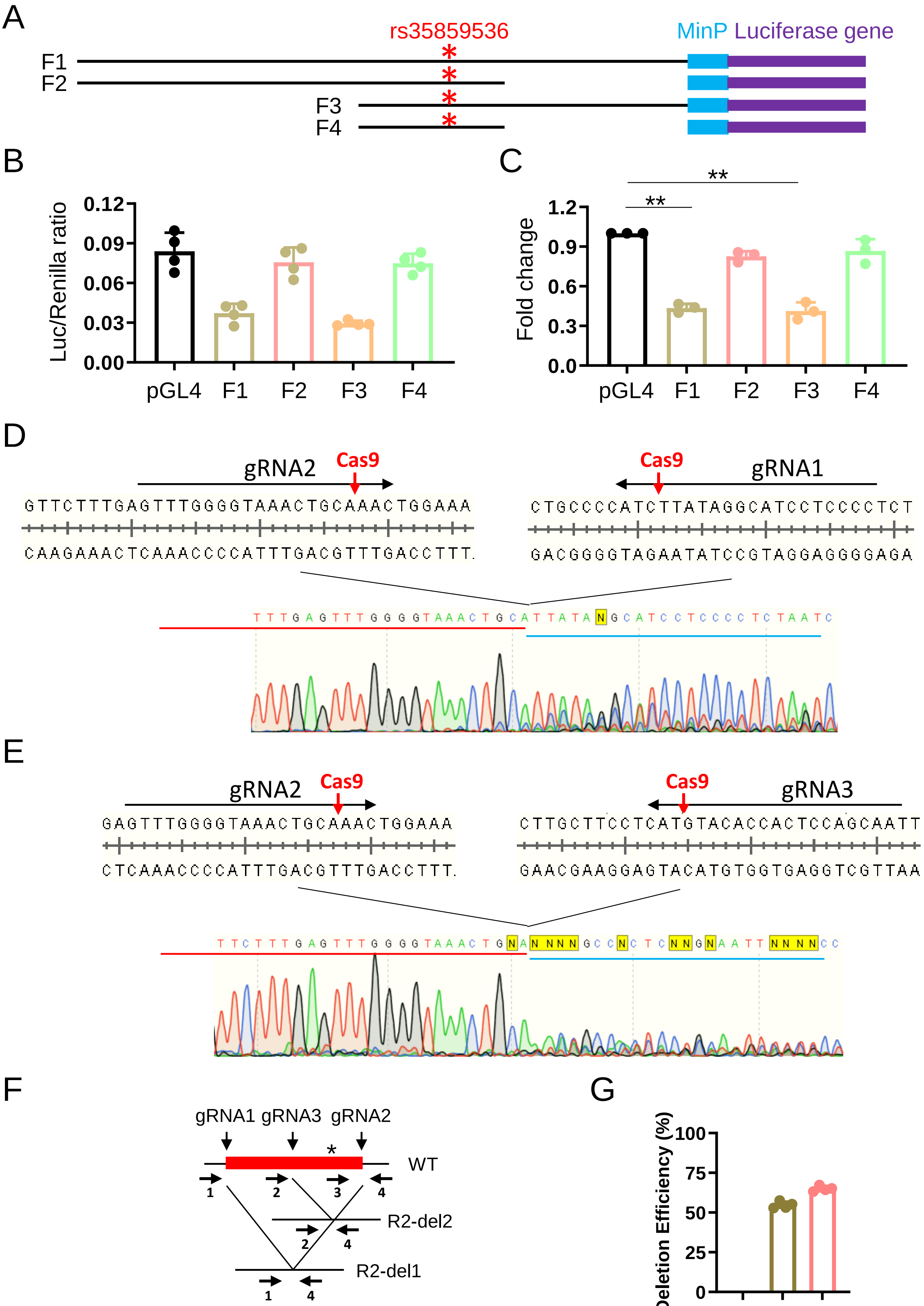

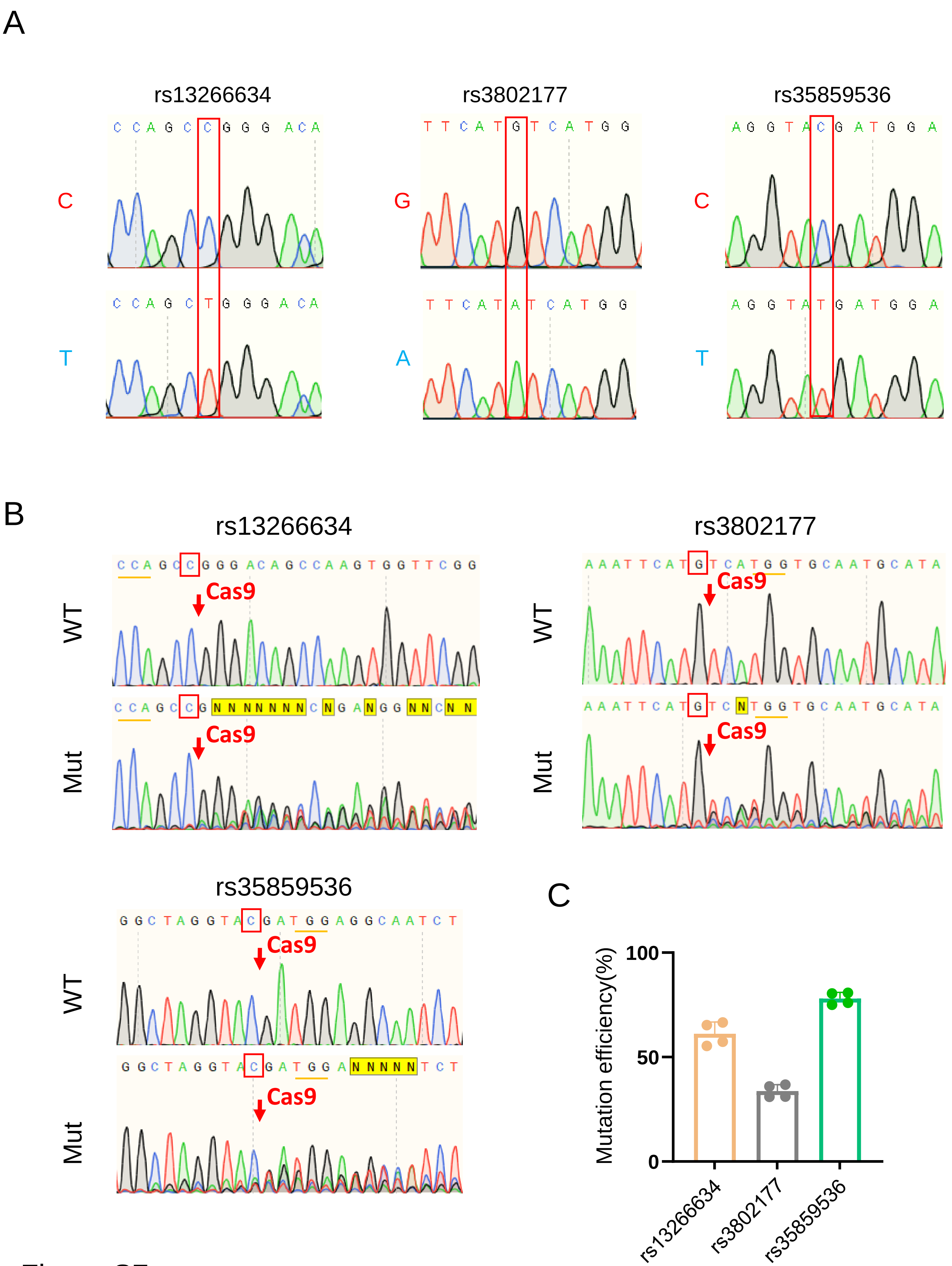

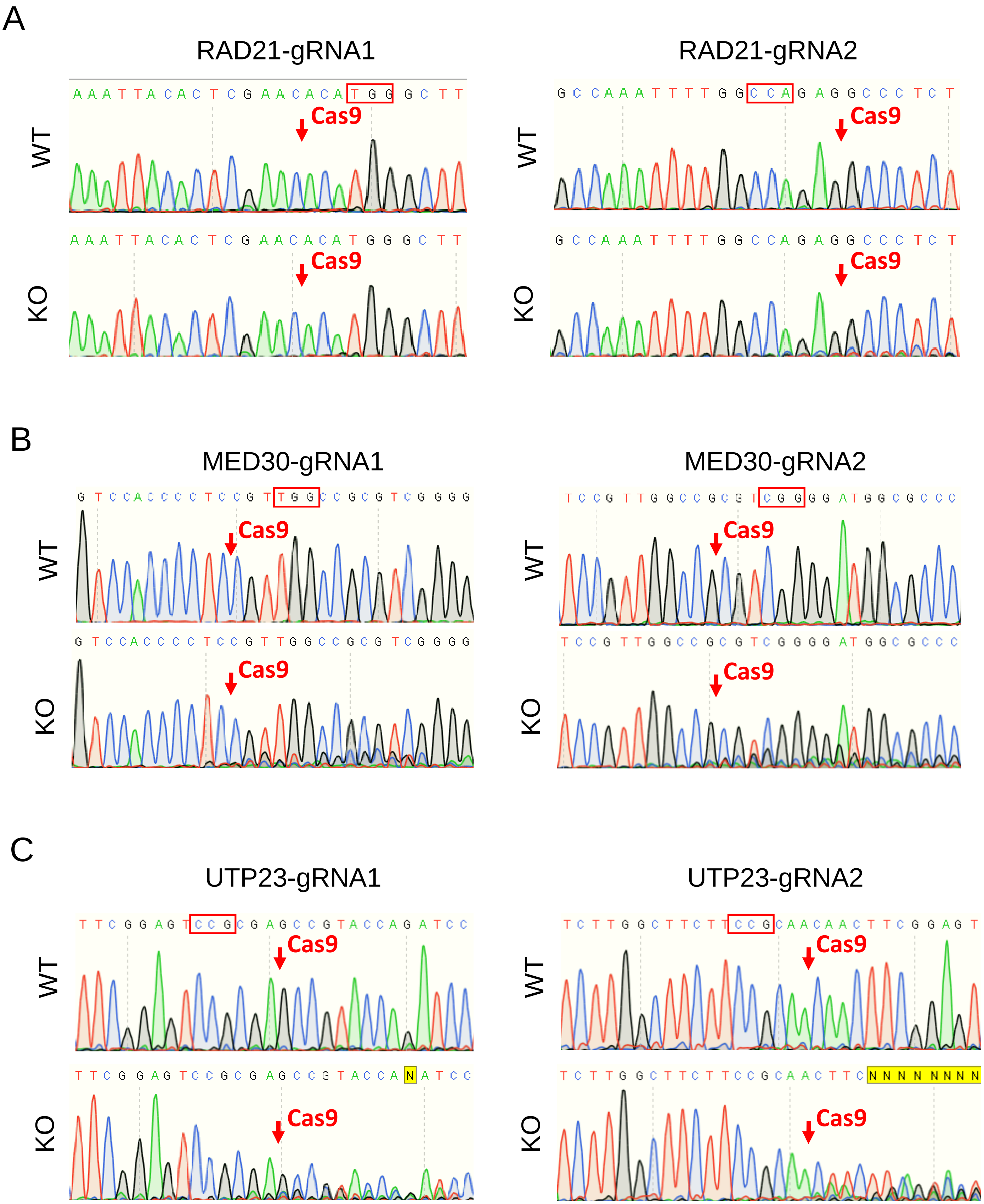

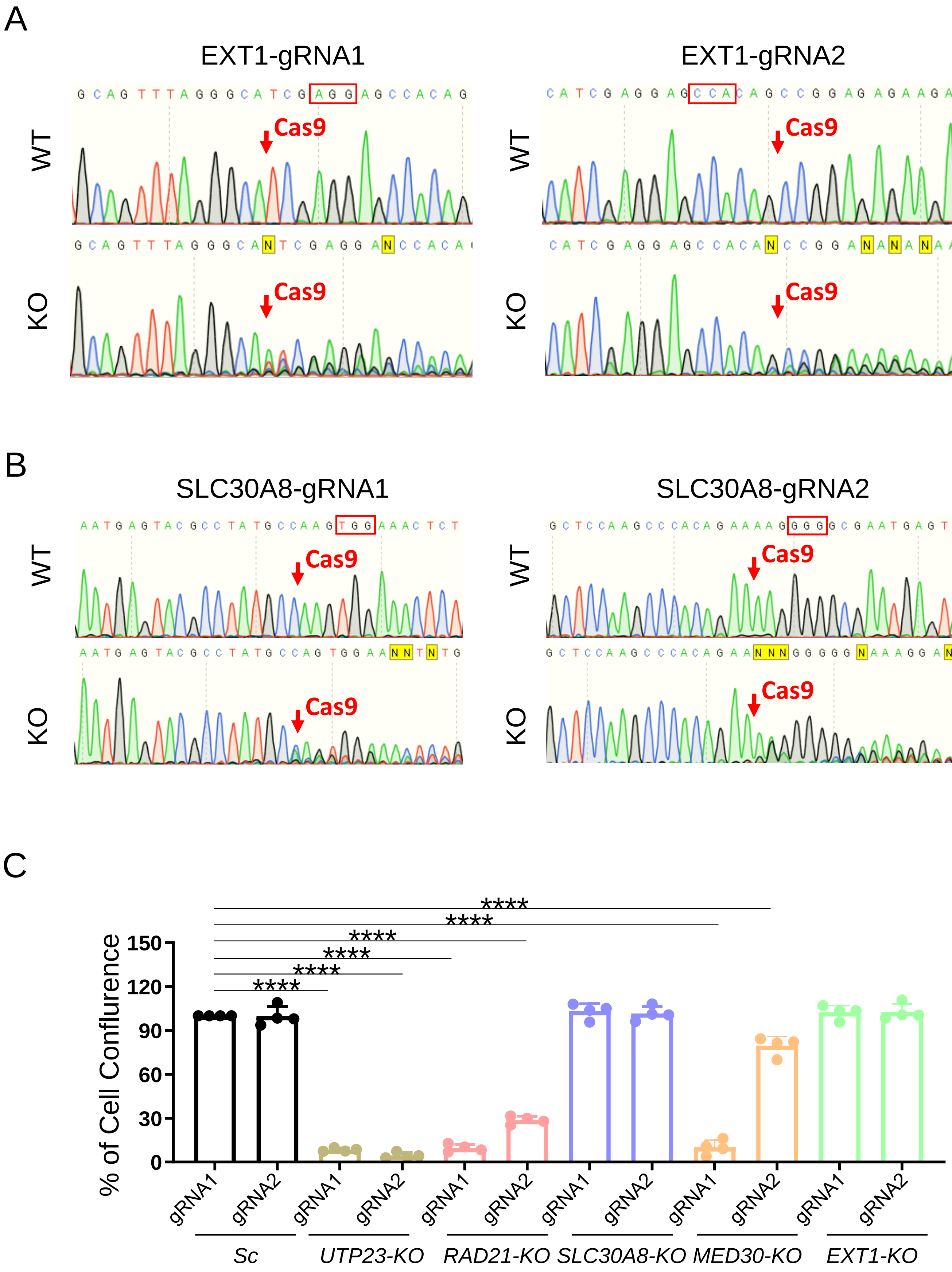

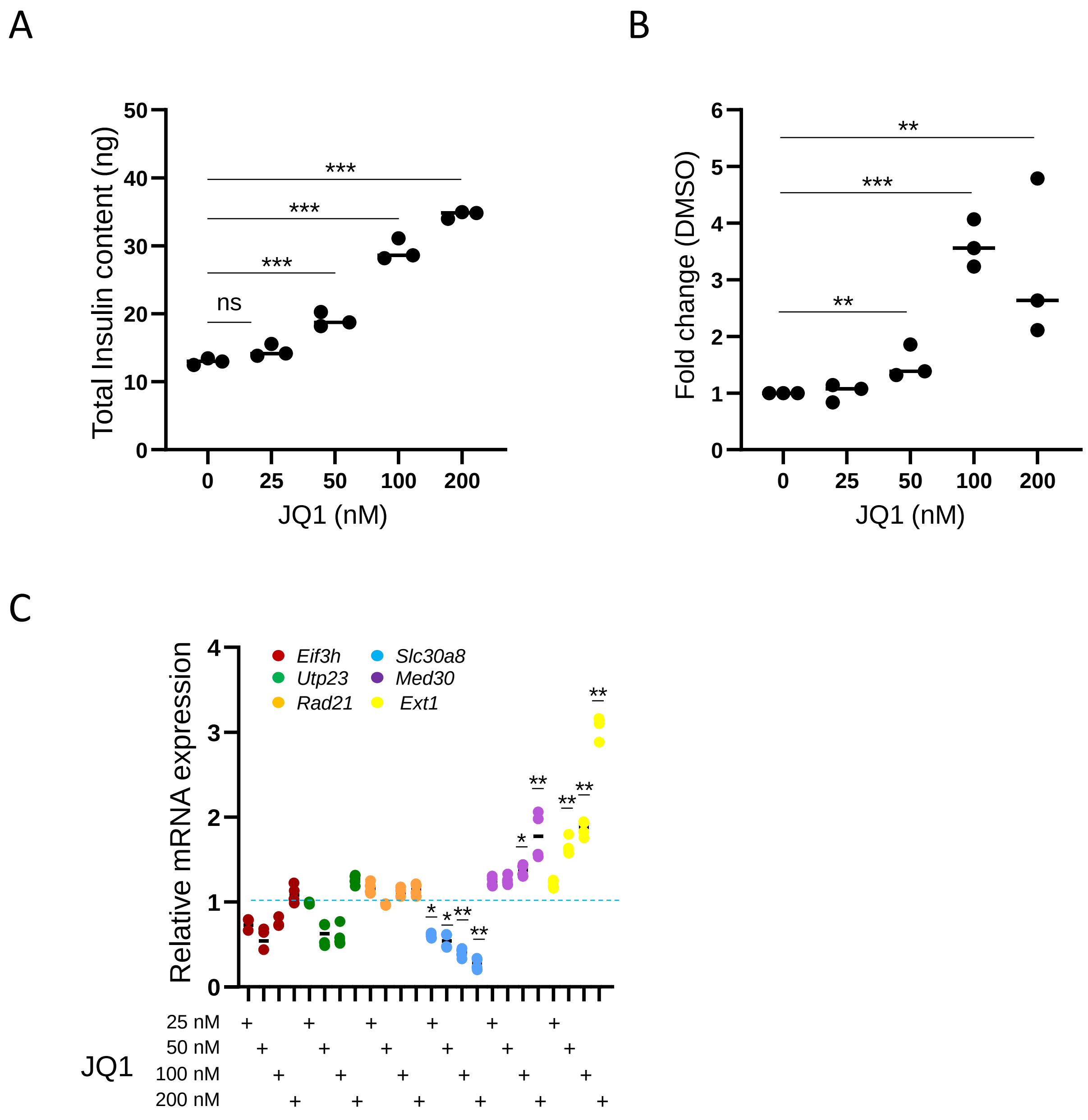

### graphical abstract

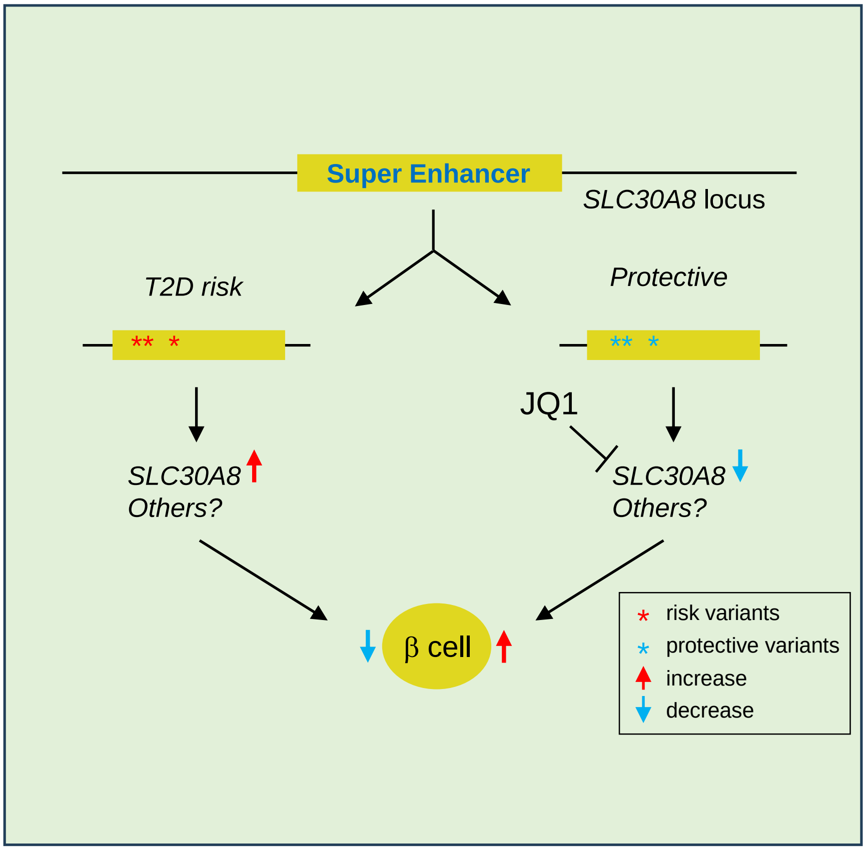
